# *Trim41* is essential for preventing X chromosome chaotic synapsis in male mice

**DOI:** 10.1101/2021.11.03.467045

**Authors:** Seiya Oura, Toshiaki Hino, Takashi Satoh, Taichi Noda, Takayuki Koyano, Ayako Isotani, Makoto Matsuyama, Shizuo Akira, Kei-ichiro Ishiguro, Masahito Ikawa

**Author notes:** **Corresponding Author**: Masahito Ikawa, Research Institute for Microbial Diseases, Osaka University, Osaka, Japan.

## Abstract

Meiosis is a hallmark event in germ cell development that accompanies sequential chromosome events executed by numerous molecules. Therefore, characterization of these factors is one of the best strategies to clarify the mechanism of meiosis. Here, we report tripartite motif-containing 41 (TRIM41), a ubiquitin ligase E3, as an essential factor for proper meiotic progression and fertility in male mice. *Trim41* KO spermatocytes exhibited synaptonemal complex protein 3 (SYCP3) overloading, especially on the X chromosome, showing extensive self-synapsis of X chromosome and non-homologous synapsis between the X chromosome and autosomes. Furthermore, the mutant mice lacking the RING domain of TRIM41, required for the ubiquitin ligase E3 activity, phenocopied *Trim41* KO mice. We then examined the behavior of mutant TRIM41 (ΔRING-TRIM41) and found that ΔRING-TRIM41 accumulated on the chromosome axes with overloaded SYCP3. This result showed that TRIM41 exerts the function on the chromosome axes. In summary, our study revealed that *Trim41* is essential for preventing SYCP3 overloading and chaotic synapsis of the X chromosome, suggesting a TRIM41-mediated mechanism for regulating unsyapsed axes during male meiotic progression.

**Summary statement:** *Trim41*-disruption caused abnormal synapsis configuration of the X chromosome and complete infertility in male mice. Thus, TRIM41 prevents the sex chromosome from chaotic synapsis.

## Introduction

Tripartite motif (TRIM) family proteins consist of more than 70 members in humans and mice (Ozato et al., 2008; Rajsbaum et al., 2014; Versteeg et al., 2013) (also according to NCBI and MGI databases). They contain three zinc-binding domains in the N-terminus: a RING finger domain, one or two B-Boxes (B1/B2) motifs, and a coiled-coil (CC) region (Fig 1A). The RING domains of many TRIM family members have been shown to confer E3 ubiquitin ligase activity (Hatakeyama, 2017; Rajsbaum et al., 2014) (Fig 1B) and implicated in various biological functions such as innate immunity and carcinogenesis regulation (Lassot et al., 2018; Zhang et al., 2013; Zhang et al., 2021). TRIM41-mediated ubiquitination also functions in natural immunity via target protein degradation, including B cell leukemia/lymphoma 10 (BCL10) (Yu et al., 2021), nucleoproteins of the influenza A virus, and vesicular stomatitis virus (Patil et al., 2020; Patil et al., 2018). Therefore, to examine the defects in natural immunity, we produced *Trim41* knockout (KO) mice. However, unexpectedly, *Trim41* KO male mice exhibited complete infertility due to meiotic defects. Thus, we focused on the analysis of meiotic events in this report.

**Fig. 1:**
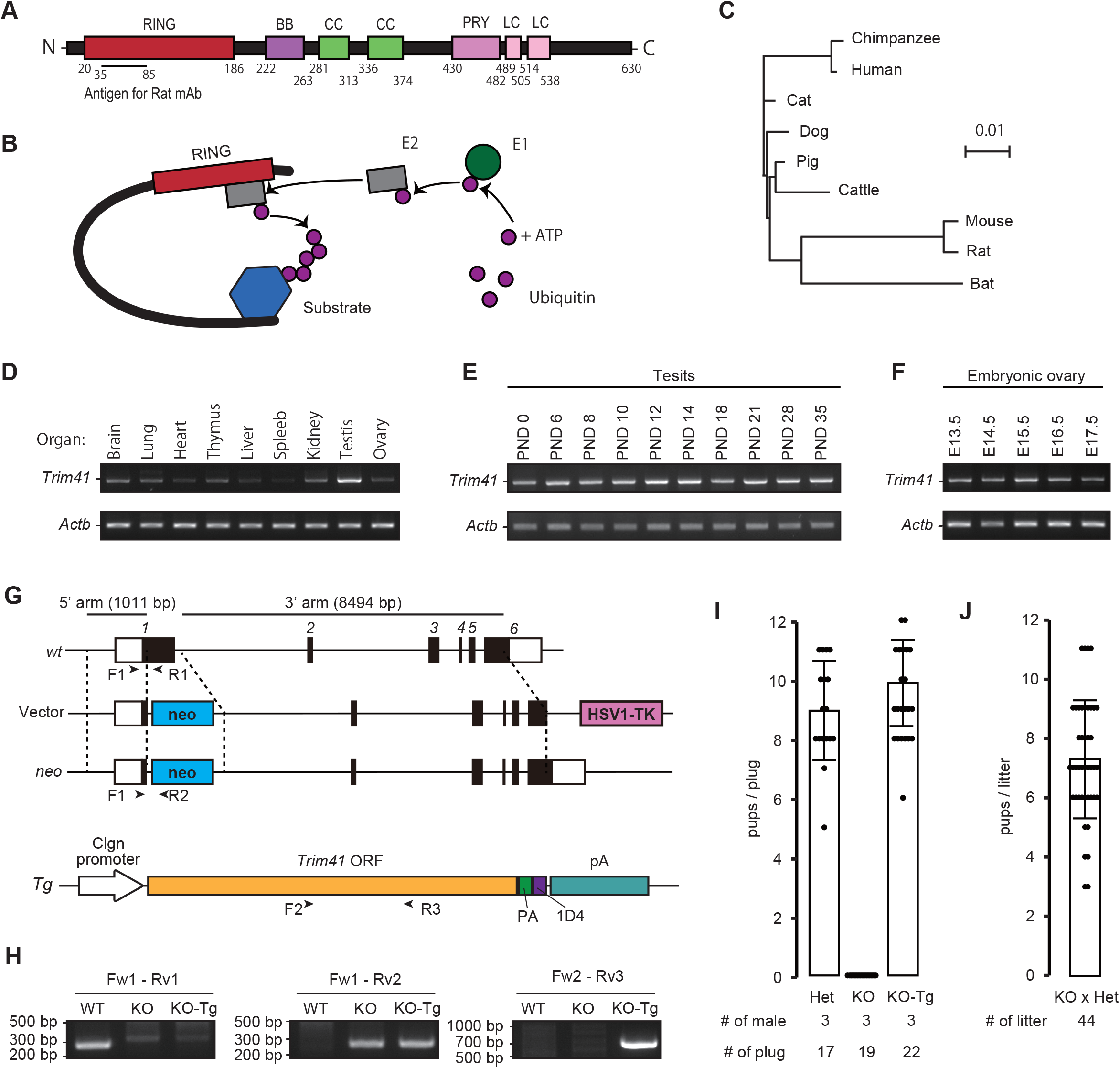
Production of *Trim41* KO mice and fertility analysis. (A) Schematic of TRIM41 protein structure and antigen position. (B) Model of TRIM E3 ubiquitin ligase activity. The RING domain at the N terminus interacts with E2 ubiquitin-conjugating enzyme. The C terminal part of TRIM recognizes the substrates. (C) Phylogenetic tree constructed by ClustalW with TRIM41 sequences of various mammals. (D) RT-PCR using multi-tissue cDNA. *Actb* was used as a loading control. (E) RT-PCR using postnatal testis cDNA. (F) RT-PCR using embryonic ovary cDNA. (G) Targeting scheme for *Trim41* disruption and a Tg contract. Black and white boxes in the gene map indicate coding and non-coding regions, respectively. Black arrowheads (Primer F1, F2, R1, R2, and R3) indicate primers for genotyping. (H) An example of genotyping PCR with primer sets in G. (I) The result of mating tests. Pups/plug: 8.9±1.7 [Het]; 0 [KO]; 9.4±2 [KO-Tg] (s.d.). Error bars indicate standard deviation. The numerical data is available in Table S3. (J) Pup numbers from mating pairs of *Trim41* KO females and *Trim41* Het males (7.4±2.3; s.d.). Error bars indicate standard deviation. The numerical data is available in Table S3.

Here, meiosis is a hallmark event in germ cell development when homologous chromosomes undergo the physical juxtaposition (called synapsis) and shuffling of genetic material via a process known as homologous recombination. Upon entry into meiosis, in the leptotene stage, programmed DNA double-strand breaks (DSBs) are introduced (Keeney et al., 2014) and enable genome-wide search for homology through the repair process, although a significant amount of chromosome pairing occurs before the initiation of DSBs (Boateng et al., 2013). This homologous search drives the pairing and alignment, finally leading to the synapsis of all homologs in the pachytene stage. However, unlike autosomes that fully synapse between homologs, sex chromosomes (X and Y in most mammals) have only synapsed at the pseudoautosomal region (PAR) and sequestered into a physically separated nuclear territory known as the XY body. The molecules involved in these complex chromosome events are not fully characterized yet, especially in mammals, due to difficulties in culturing and genetically manipulating spermatogenic cells *in vitro*. Thus, physiological screening with KO mice has been a powerful strategy to identify meiosis-related factors (Gray and Cohen, 2016; Oji et al., 2020; Oura et al., 2021).

In this study, we revealed that *Trim41* deficiency cause synaptonemal complex protein 3 (SYCP3) overloading on several chromosome axes. Among them, the X chromosome was mainly affected, exhibiting aberrant synapsis with autosomes rather than forming the XY body. A transgene with a germ cell-specific expression rescued the spermatogenesis defects, which showed that TRIM41 directly regulates meiosis. Further, deletion of the RING domain (ΔRING) phenocopied *Trim41* KO. More importantly, ΔRING-TRIM41 accumulated on chromosome axes with overloaded SYCP3, suggesting that TRIM41-mediated protein degradation may act for removing overloaded SYCP3. Thus, our study demonstrated that mammalian spermatocytes have a TRIM41-mediated mechanism for preventing sex chromosome chaotic synapsis.

## Results

### *Trim41* is evolutionarily conserved between mammals and highly expressed in pachytene germ cells

The Treefam database (http://www.treefam.org/; Release 9, March 2013) shows that TF342569, a family containing TRIM41, has been annotated in 50% of eukaryotes, 98 % of vertebrates, and 100% of mammals (Fig. S1A). Also, phylogenetic analysis with Clustal W2.1(Larkin et al., 2007) showed that TRIM41 was highly conserved in many mammals, including cattle, dogs, mice, and humans (Figs 1C and S1B). Next, to determine the expression profile of *Trim41*, we performed RT-PCR using cDNA obtained from adult tissues, postnatal testis, and embryonic ovary. These RT-PCR showed that *Trim41* is expressed ubiquitously (Figs 1D–1F), albeit the most highly in testis (Fig. 1D). Furthermore, according to published single-cell RNA-sequencing (scRNA-seq) data analyzing mouse testis (Hermann et al., 2018) and embryonic ovary (Niu and Spradling, 2020), *Trim41* was the most highly expressed in pachytene germ cells (Figs S1C and S1D). These results suggest that TRIM41 functions during the meiotic phase of mammalian gametogenesis.

### *Trim41* is essential for male fertility

To uncover the function of *Trim41 in vivo*, we generated *Trim41* KO mice by transfecting embryonic stem cells (ESCs) with a targeting vector (Fig. 1G; vector). Targeted ESC clones were injected into ICR embryos to obtain chimeric mice. The chimeric mice were mated with wild-type (WT) females to establish *Trim41* KO lines (Fig. 1H). KO mice obtained by heterozygous intercrosses showed no overt gross defects in development, behavior, and survival. Then, we housed individual *Trim41* KO male mice with wild type (wt) females for two months to analyze their fertility. Although we observed 19 vaginal plugs, *Trim41* KO males failed to sire pups (Fig. 1I). On the other hand, we obtained pups from the mating pair of *Trim41* KO females and *Trim41* Het males (7.4±2.3; Fig. 1J), indicating that *Trim41* is dispensable for female fertility. As *Trim41* Het male mice are fully fertile, we used littermate heterozygous males as controls in some experiments.

### A transgene under *Clgn* promoter restored infertility of *Trim41* KO males

To confirm infertility phenotype of *Trim41* KO males was due to *Trim41* deficiency, not an aberrant genetic modification near the *Trim41* locus, and rule out the possibility that latent systemic abnormalities affected male fertility, we carried out a tissue-specific rescue experiment by generating transgenic (Tg) mouse lines. First, we injected a DNA construct having PA-1D4-tagged *Trim41* under the testis-specific Clgn promoter (Watanabe et al., 1995) (Fig. 1G; Tg) and established a Tg line (Fig. 1H; Fw2-Rv3). *Clgn* promoter-driven *Trim41* expression started around postnatal day (PND) 9–11 (Fig. S2A), corresponding to the spermatocyte appearance. The Tg expression was also confirmed by immunoblotting analysis using anti-PA (Fig. S2B) and -TRIM41 antibody (Fig. S2C), although the expression level of Tg was lower than that of intrinsic *Trim41* (Fig S2C; WT v.s. KO-Tg). Then we housed *Trim41* KO male mice expressing the Tg (referred to as KO-Tg) with WT females and observed the normal count of pups (Fig. 1I), showing the *Clgn* promoter-driven *Trim41* Tg rescued the fertility of KO males. These results indicated that *Trim41* expression from meiotic entry onward in testis is essential for male fertility.

### *Trim41* KO male mice exhibited oligozoospermia

When we observed gross testis morphology, *Trim41* KO testes were smaller than those of control (testis/ body weight: 3.2±0.3 x 10^−3^ [WT], 3.2±0.2 x 10^−3^ [Het],1.2±0.2 x 10^−3^ [KO]; Figs 2A and 2B), indicating defective spermatogenesis in *Trim41* KO testis. To define the cause of testicular atrophy, we performed hematoxylin and periodic acid-Schiff (HePAS) staining of testicular sections. While three germ cell layers existed in control testis sections, only two layers of germ cells existed in *Trim41* KO testis (Fig. 2C; low magnification). The number of spermatocytes and spermatids dramatically decreased in the KO testis (Fig. 2C; stage VII–VIII and XII). Although a few elongating/elongated spermatids existed in the KO testis, their nuclei were not fully compacted (Fig. 2C; red arrows). Consistent with the dramatic decrease of spermatocytes and spermatids, the number of TUNEL positive cells increased in KO testis compared with Het counterparts (Figs 2D and 2E). The TUNEL positive cells were frequently observed in the second layer (Fig. 2D; high magnification). We did not see the accumulation of TUNEL positive cells in a specific stage of the seminiferous epithelium cycle (Fig. 2E). These results showed that spermatocytes after the pachytene stage were gradually eliminated by apoptosis. As a result, only a few mature spermatozoa existed in the cauda epididymis (Fig. 2F). In addition, all the spermatozoa exhibited abnormal head/tail shapes (Fig. 2G) and were immotile. These *Trim41* KO spermatogenesis defects were restored by *Clgn-Trim41* (Figs 2A–2F), albeit partially in TUNEL analysis (Fig 2F). These observations suggested that *Trim41* KO males exhibited spermatogenesis defects, leading to oligozoospermia.

**Fig. 2:**
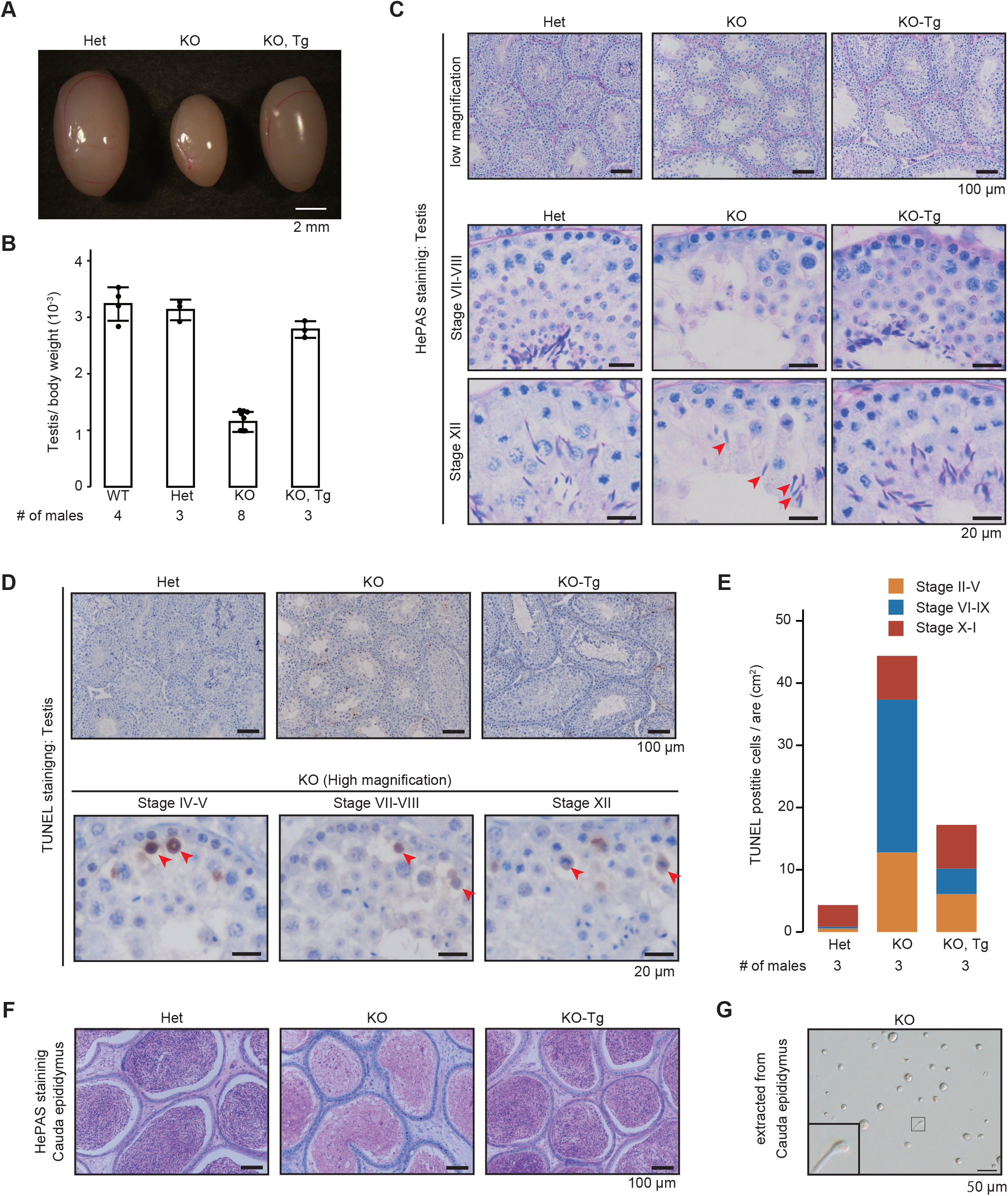
Histological analysis of *Trim41* KO male mice. (A and B) Testis morphology (A) and testis/bodyweight of WT, *Trim41* Het, *Trim41* KO, and KO-Tg adult mice (B). Testis/body weight: 3.2±0.3 x 10–3 [WT], 3.1±0.2 x 10–3 [Het], 1.2±0.2 x 10^−3^ [KO], 2.8±0.1 x 10^−3^ [KO-Tg]. Error bars indicate standard deviation. The numerical data is available in Table S3. (C) PAS staining of seminiferous tubules of adult mice. The seminiferous epithelium cycle was determined by germ cell position and nuclear morphology. Red arrows indicate elongated spermatids with abnormal head morphology. At least three male mice were analyzed. (D) TUNEL staining of seminiferous tubules of adult mice counterstained with hematoxylin. At least three male mice were analyzed. (E) The stacked bar graph of TUNEL positive cells. The seminiferous epithelial stages were roughly determined by the arrangement and nuclear morphology of the first layer of germ cells (spermatogonia and leptotene/zygotene spermatocytes). The numerical data is available in Table S3. (F) PAS staining of cauda epididymis of adult mice. At least three male mice were analyzed. (G) Spermatozoa extracted from cauda epididymis of adult mice.

### *Trim41* KO spermatocytes underwent SYCP3 overloading

Due to an apparent defect during meiosis, we examined DNA DSBs and synapsis by immunostaining surface chromosome spreads with anti-γH2AX and SYCP3, respectively (Figs 3A and 3B). The SYCP3/γH2AX immunostaining pattern in leptotene and zygotene spermatocytes was comparable between the two genotypes, showing that *Trim41* KO spermatocytes underwent programmed DSBs and initial assembly of the synaptonemal complex. However, the compaction of the XY axes, a telltale signature of XY body formation, was rarely observed in KO spermatocytes (Figs 3B and 3C). Consistent with the XY body malformation, the pachytene and diplotene populations decreased dramatically in KO testis (Fig. 3D). More strikingly, intense SYCP3 signals were observed on both autosomes and sex chromosomes (Figs 3B and 3C). The autosomes with the overloaded-SCYP3 tended to be inside of γH2AX signals, suggesting that the overloading of SCYP3 correlates with autosome unsynapsis. Furthermore, we also observed multilayer SCYP3 axes in the X chromosome (Fig. 3C; yellow arrowhead) and juxtaposition of sex chromosome and autosomes (Fig. 3C; red arrowhead). Such overloading of SYCP3 became evident once homolog synapsis had completed at pachytene rather than at zygotene/pachytene transition (zygotene/pachytene transition; Figs S3A and S3B).

**Fig. 3:**
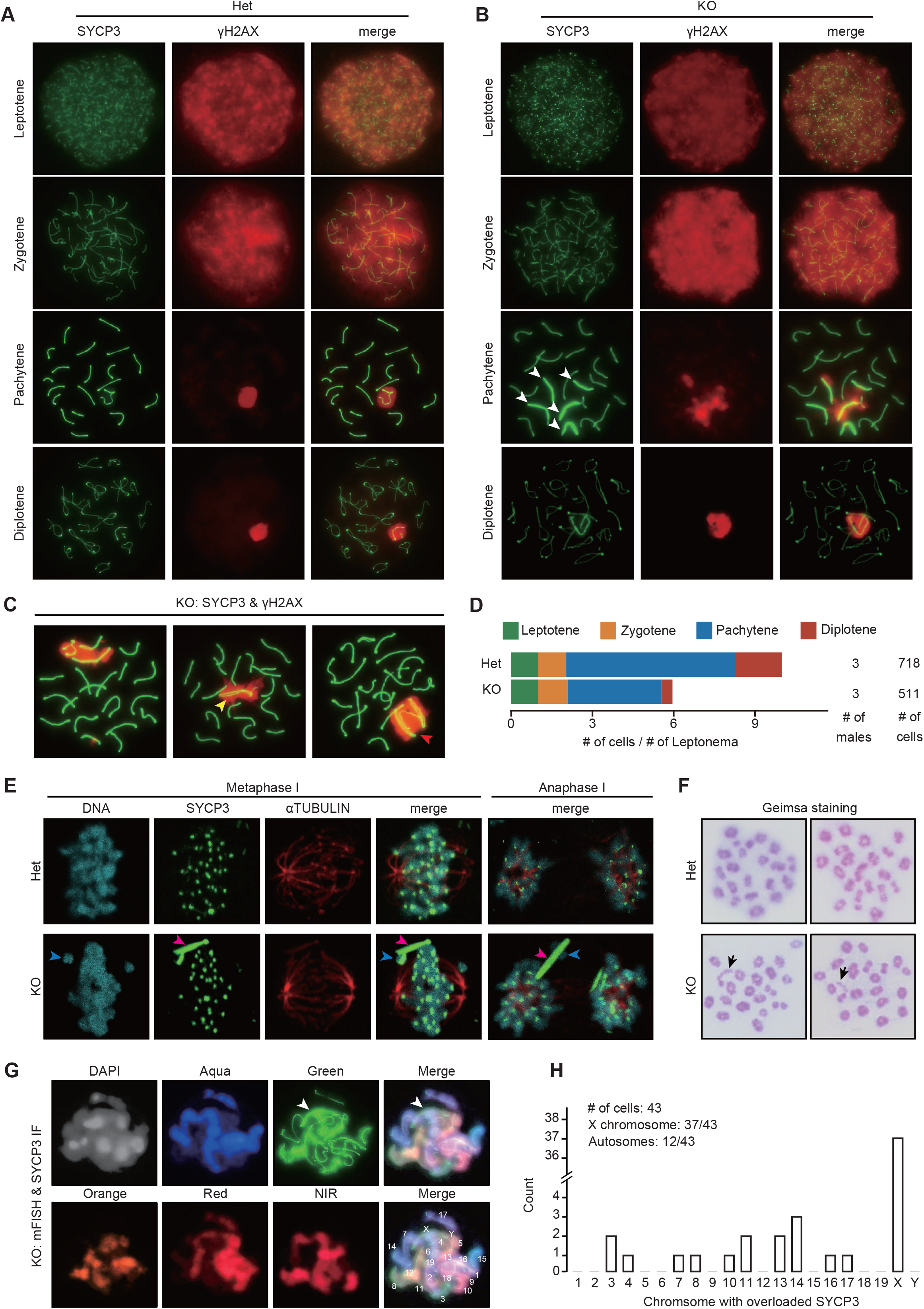
Cytological analysis of *Trim41* KO spermatocytes. (A and B) Immunostaining of spread nuclei from prophase spermatocytes collected from adult Het (A) and KO (B) male mice. White arrows indicate overloaded SYCP3 axes. At least three male mice were analyzed. (C) Additional immunostaining images of KO spermatocytes. The yellow arrowhead indicates the X chromosome with two SYCP3 axes. The red arrowhead shows the evident synapsis between the sex chromosome and autosomes (only 17 axes existed outside the γH2AX cloud). (D) The percentage of each meiotic prophase stage in immunostained spread nuclei samples in A and B. The numerical data is available in Table S3. (E) Immunostaining of metaphase and anaphase spermatocytes squashed from seminiferous tubules after fixation. Magenta and blue arrowheads indicate polycomplex-like structures, aggregate of SYCP3, and chromosomes connected with the polycomplex-like structures. At least three male mice were analyzed. (F) Giemsa staining of spread nuclei of metaphase I spermatocytes. Black arrows indicate the miscondensation of chromosomes. At least three male mice were analyzed. (G) Multicolor FISH followed by SYCP3 immunostaining. White arrows indicate overloaded SYCP3 axes. (H) The histogram for the chromosome distribution of SYCP3 overloaded axes. Forty-three well-spread cells were analyzed.

Since localization of axial element proteins, SYCP2 and SYCP3, depends on cohesin axial core that is generated by REC8 and RAD21L (Fujiwara et al., 2020; Ishiguro et al., 2014; Ishiguro, 2019), we examined the REC8 and RAD21L in *Trim41* KO spermatocytes. Consistent with SYCP3 overloading, REC8 and RAD21L tended to be hyperaccumulated (Fig. S4). Although RAD21L gradually dissociates from axial elements from late pachytene onwards in WT spermatocytes (Fig. S4C) (Ishiguro et al., 2011), the dissociation was evident in early pachytene of KO spermatocytes (Fig. S4D). This observation is at least in part consistent with the evidence that Trim41 is co-immunoprecipitated by RAD21L (Fujiwara et al., 2020).

Interestingly, the overloaded SYCP3 remained on a few chromosomes even in metaphase I and anaphase I (Fig. 3E; blue arrowheads). Further polycomplex-like structure, aggregate of SYCP3, was also observed apart from chromosomes (Fig. 3E; magenta arrowheads). In addition, some chromosomes connected with the SYCP3 failed to align in the equatorial plate (Fig. 3E; Metaphase I) and were left behind between two centrosomes (Fig. 3E; Anaphase I). Probably because of the remained SYCP3 axes, one or two chromosomes failed to form bivalents and remained as univalents at metaphase I (Fig. 3F). Because chromosome alignment requires a tension across homologs during metaphase I (Kim et al., 2015; Woods et al., 1999), it was likely that lack of chiasmata caused misalignment of those univalents. These results showed that the SYCP3 amount on chromosome axes and synapsis configuration was misregulated in *Trim41* KO spermatocytes.

### SYCP3 overloading were biased to the X chromosome

Then, to examine whether SYCP3 overloading is biased to specific chromosomes, we combined SYCP3 immunostaining with multicolor fluorescence in situ hybridization (mFISH; Fig. 3G). The X chromosomes showed more frequent SYCP3 overloading than autosomes (Fig. 3H; X chromosome: 37 out 43 cells; autosome: 12 out 43 cells). Of note, the autosomes with overloaded SYCP3 signals were almost random (Fig. 3H). Finally, we examined the pachytene stage in the E15.5 female ovary. The gross gonad morphology was comparable between the two genotypes (Fig. S5A). The SYCP3 overloaded axes were observed only in a few germ cells (Fig. S5B). This result was contrast to the observation that SYCP3 overloading was frequent on X chromosome in spermatocytes (Fig. 3H). As a result, folliculogenesis progressed normally (Fig. S5C), and *Trim41* KO female mice were fully fertile (Fig. 1M). Overall, these results showed that *Trim41* is essential for the proper behavior of the unsynapsed X chromosome.

### SYCP3 overloaded X chromosome saw self- and non-homologous-synapsis

Next, to examine the synapsis state, we visualized the central element of the synaptonemal complex (synapsed axes; Figs 4A and 4B) and unsynapsed axes (Figs 4C and 4D) by immunostaining surface chromosome spreads with anti-SYCP1 and BRCA1 antibodies, respectively. SYCP1 is a molecule known to localize between the lateral element of the synaptonemal complex of homologous chromosomes and form transverse filaments, rungs of a ladder of synapsis (Liu et al., 1996; Meuwissen et al., 1992). BRCA1, the product of the breast cancer 1, accumulates on unsyapsed axes of sex chromosomes (Scully et al., 1997). Consistent with Fig. 3C (indicated by a red arrow), X chromosome and autosome were connected by SYCP1 (Fig. 4B; white arrow) in some spermatocytes. SYCP1 signals tended to remain on the overloaded SYCP3 of X chromosomes (Fig. 4B), indicating that the X chromosomes synapse by themselves or between sister chromatids. Strikingly, we observed multilayered SCYP1 signals in some spermatocytes (Fig. 4B; blue arrows).

**Fig. 4:**
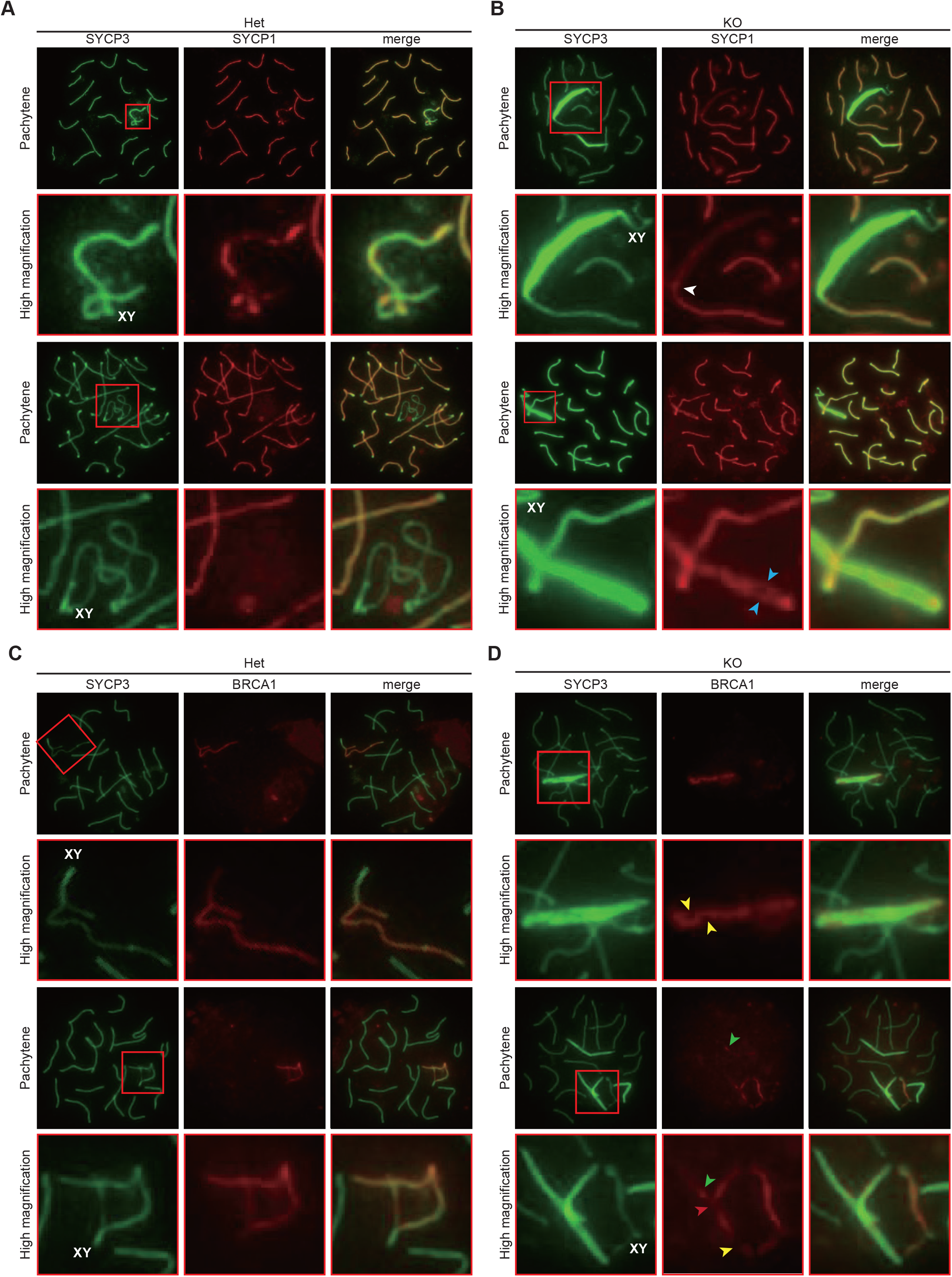
Synapsis configuration of SYCP3 overloaded axes. (A and B) SYCP1 immunostaining of spread nuclei from prophase spermatocytes collected from adult Het (A) and KO (B) male mice. The sex chromosomes were identified by their characteristic shape of SYCP3 axes. At least three male mice were analyzed. (C and D) BRCA1 immunostaining of spread nuclei from prophase spermatocytes collected from adult Het (C) and KO (D) male mice. At least three male mice were analyzed.

On the other hand, the BRCA1 staining pattern became uneven throughout the X chromosomes (Fig. 4D; yellow and red arrows), indicating that the X chromosomes have synapsed somehow. Furthermore, BRCA1 was positive on several autosomes with overloaded SYCP3, suggesting that meiotic silencing of unsynapsed chromatin (MUSC) was activated presumably as a result of delayed homolog synapsis at the SYCP3-overladed region. (Fig. 4D; green arrows). More notably, we observed the BRCA1 negative interface between the X chromosome and autosome axes (Fig. 4D; red arrow), suggesting that the X chromosome synapses with the autosome. These results indicated the X chromosome synapsed by themselves or with autosomes in *Trim41* KO spermatocytes.

### *Trim41^ΔRING/ΔRING^* male mice phenocopied *Trim41^neo/neo^* male mice

To examine whether the RING domain is essential for *Trim41* function, we disrupted the RING domain by inserting the HA affinity tag sequence (Fig. 5A; referred to as ΔRING). The ΔRING-TRIM41 loses only ubiquitin ligase activity, not interaction with target proteins (Patil et al., 2020; Patil et al., 2018). We microinjected two gRNA/Cas9 ribonucleoprotein complexes and a ssODN into zygotes (Fig. 5A). Of 145 injected eggs, 119 eggs reached the two-cell stage. Then we transplanted the two-cell eggs into the oviducts of six pseudopregnant female mice and obtained twenty-five pups. The genotype PCR screening (Fig. 5B) detected the intended mutation with a 180 bp deletion and HA tag insertion (referred to as *Trim41^ΔRING^*) from ten pups. Immunoblotting analysis with anti-HA and -TRIM41 antibodies detected HA-tagged ΔRING-TRIM41 expression (Figs 5C and 5D). Then, we caged *Trim41^ΔRING/ΔRING^* (referred to as ΔR-Homo; *Trim41^wt/ΔRING^*: referred to ΔR-Het) male mice with WT females for two months to analyze their fertility. Although we confirmed 38 vaginal plugs, *Trim41* ΔR-Homo males failed to sire any pups (Fig. 5E) as well as *Trim41* KO males. Further, testis of *Trim41* ΔR-Homo males was smaller than the ΔR-Het counterparts (Figs 5F and 5G) due to spermatogenesis defects (Fig. 5H). The TUNEL-positive cells were frequently observed in the 2nd layer of tubules (Fig. 5I). Also, TUNEL signals were positive in overall seminiferous epithelial cycles (Fig. 5J). As a result, almost no fully-matured spermatids existed in the cauda epididymis of ΔR-Homo males (Fig. 5K). These results showed that *Trim41^ΔRING/ΔRING^* male mice exhibited the same phenotype as *Trim41^neo/neo^* male mice, suggesting that the RING domain is essential for TRIM41 function.

**Fig. 5:**
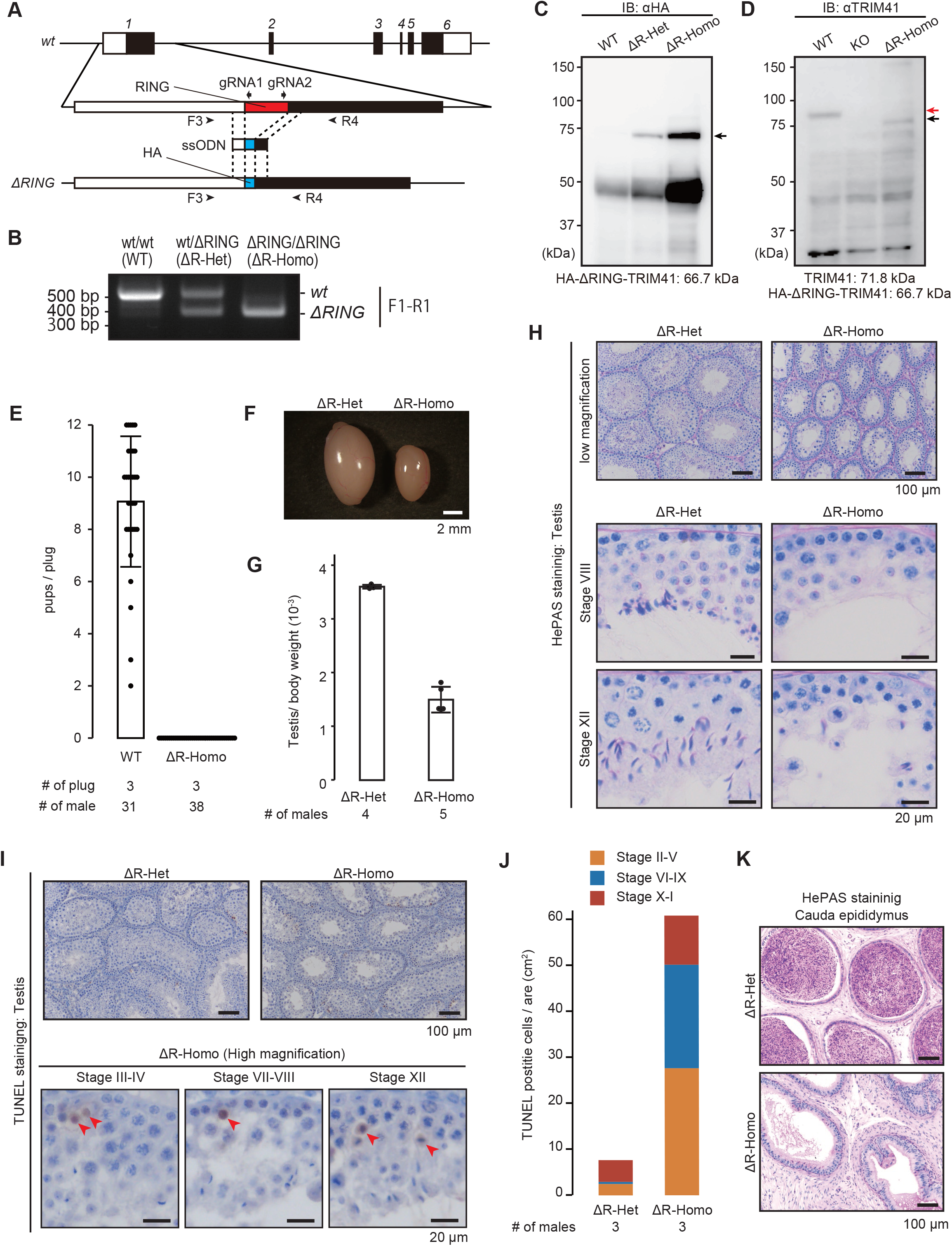
Production and phenotypical analysis of *Trim41^ΔRING/ΔRING^* mice. (A) Knock-in scheme for the replacement of the RING domain with HA tag. Black and white boxes in the gene map indicate coding and non-coding regions, respectively. Black arrowheads (F3 and R4) and arrows (gRNA1 and gRNA2) indicate primers for genotyping and target sequence of gRNAs, respectively. (B) An example of genotyping PCR with primer sets in A. (C and D) Immunoblotting analysis with anti-HA (C) and -TRIM41 (D) antibodies. Black and red arrows indicate TRIM41 and HA-ΔRING-TRIM41, respectively. (E) The result of mating tests. Pups/plug: 9.1±2.5 [WT]; 0 [ΔR-Homo] (s.d.). Error bars indicate standard deviation. The numerical data is available in Table S3. (F and G) Testis morphology (F) and testis/bodyweight of ΔR-Het and ΔR-Homo adult mice (G). Testis/body weight: 3.6±0.0 x 10^−3^ [ΔR-Het]; 1.5±0.2 x 10^−3^ [ΔR-Homo]. Error bars indicate standard deviation. The numerical data is available in Table S3. (H) PAS staining of seminiferous tubules of adult mice. The seminiferous epithelium cycle was determined by germ cell position and nuclear morphology. At least three male mice were analyzed. (I) TUNEL staining of seminiferous tubules of adult mice counterstained with hematoxylin. At least three male mice were analyzed. (J) The stacked bar graph of TUNEL positive cells. The seminiferous epithelial stages were roughly determined by the arrangement and nuclear morphology of the first layer of germ cells (spermatogonia and leptotene/zygotene spermatocytes). The numerical data is available in Table S3. (K) PAS staining of cauda epididymis of adult mice. At least three male mice were analyzed.

### ΔRING-TRIM41 accumulated on SYCP3-overlaoded region

Immunostaining analysis with anti-SYCP3 and γH2AX antibodies demonstrated XY body malformation and SYCP3 overloading in ΔR-Homo pachytene spermatocytes (Figs 6A and 6B). Also, the number of pachytene and diplotene spermatocytes decreased dramatically in ΔR-Homo testis (Figs S6A–C). These results corroborated the necessity of the RING domain in TRIM41 functions.

**Fig. 6:**
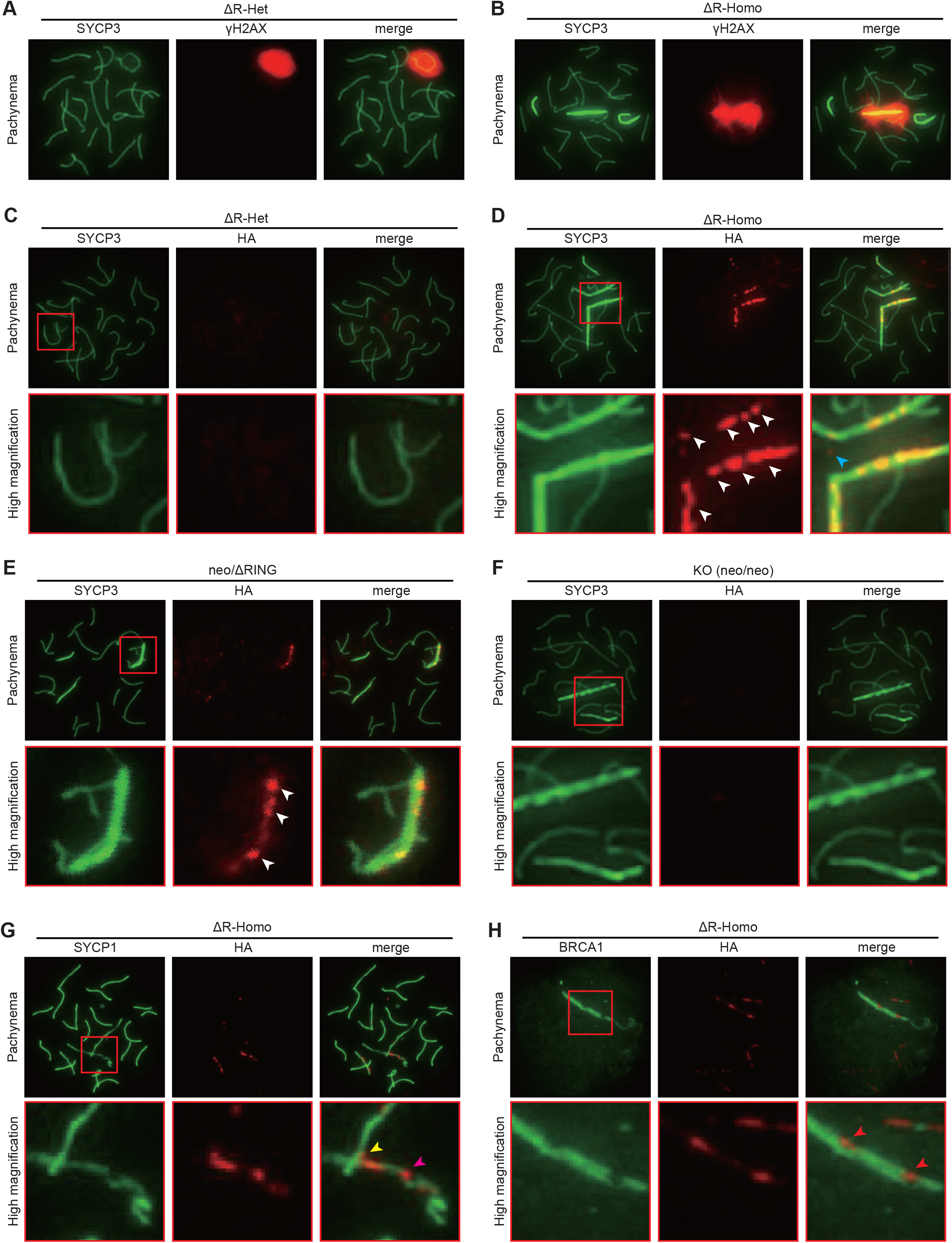
Cytological analysis of *Trim41^ΔRING/ΔRING^* spermatocytes. (A and B) SYCP3/γH2AX immunostaining of pachytene spermatocytes from ΔR-Het (A) and ΔR-Homo (B) male mice. At least three male mice were analyzed. (C–F) SYCP3/HA immunostaining of pachytene spermatocytes from ΔR-Het (C), ΔR-Homo (D), *Trim41^neo/ΔRING^* (E), and KO (F) male mice. White and blue arrowheads indicate HA signals on and outside of SYCP3 axes, respectively. At least three male mice were analyzed for ΔR-Het and ΔR-Homo. (G) SYCP1/HA immunostaining of surface chromosome spread from ΔR-Homo testis. Magenta and yellow arrowheads indicate HA signals on the SYCP1 positive/negative boundary and the non-homologous axis interface, respectively. At least three male mice were analyzed. (H) BRCA1/HA immunostaining of surface chromosome spread from ΔR-Homo testis. Red arrowheads indicate HA signals on the BRCA1 negative part of the X chromosome axes. At least three male mice were analyzed.

Next, to determine TRIM41 localization, we stained the spread nuclei with an anti-HA antibody (Figs 6C–F). In ΔR-Het spermatocytes, we observed only faint signals throughout the nuclei (Fig. 6C). However, surprisingly, ΔR-Homo spermatocytes saw intense HA signals on the SYCP3-overloaded regions (Figs 6D and S6A; white arrows). The strong HA signals appeared after the zygotene–pachytene transition (Figs S7A). In addition, the HA signals tended to become stronger during prophase I progression (Figs S7A), showing a correlation with SYCP3 overloading. The HA foci were also detected in *Trim41^neo/ΔRING^* spermatocytes (Fig. 6E), demonstrating that the removal of the RING domain changed the immunostaining pattern, not the protein level difference. On the other hand, we did not detect the HA immunostaining signals in *Trim41* KO spermatocyte with SYCP3 overloading (Fig. 6F), ruling out the possibility of antibody cross-reaction. Furthermore, we confirmed the same phenomena in immunostaining analysis with an anti-TRIM41 antibody (Figs S7B–E). These observations showed that ΔRING-TRIM41 accumulated on the SYCP3-overloaded regions, suggesting that TRIM41 functions on the chromosome axes. As a side note, we also identified several HA foci outside of the chromosome axes in zygotene and early pachytene spermatocytes (Figs 6D and S7A; blue arrows), although we could not specify the location of those foci.

### ΔRING-TRIM41 accumulated on the synapsed part of the X chromosome

Finally, we examined the relationship between ΔRING-TRIM41 accumulation and the synapsis state by SYCP1 and BRCA1 immunostaining. First, we confirmed SYCP1 tended to remain on the sex chromosome axes in ΔR-Homo spermatocytes and KO spermatocytes, although the sex chromosome SYCP1 axes were uneven (Figs. 6G and S8A). Then, we compared the SYCP1 immunostaining pattern with the ΔRING-TRIM41 accumulation pattern and noticed that the boundary of the SYCP1 positive and negative parts were hotspots of ΔRING-TRIM41 accumulation (Figs. 6G and S8A; magenta arrowheads). More strikingly, the ΔRING-TRIM41 also existed on the interface between the X chromosome and autosomes (Figs. 6G and S8A; yellow arrowheads). On the other hand, in BRCA1/HA immunostaining analysis, we observed ΔRING-TRIM41 on the BRCA1 negative part of the X chromosome axes (Figs. 6H and S8B; yellow arrowheads). Above all, these observations suggested that ΔRING-TRIM41 accumulated on the synapsed part of the X chromosome axes, and therefore TRIM41 is essential for preventing the chaotic X chromosome synapsis.

Given that TRIM41 confers ubiquitin ligase E3 activity (Patil et al., 2020; Patil et al., 2018), we tried to identify the substrates by immunoprecipitation. However, we could not detect SYCP3 in co-IPed elute samples by immunoblotting (Figs S9A–S9F). Also, we could not find candidate proteins in IP-Mass analysis (Figs S9G–S9I). Thus, substrate discovery would be the feature research.

## Discussion

In this research, we produced *Trim41* KO mice and analyzed the phenotype. Unexpectedly, *Trim41* deficiency caused meiosis defects and infertility in male mice. The infertility was rescued by a transgene driven by a testis-specific *Clgn* promoter. In detailed analyses, we found that *Trim41* KO spermatocytes exhibited SYCP3 overloading. We also noticed that and the X chromosomes underwent self- and non-homologous-synapsis. Furthermore, we removed the RING domain of TRIM41 (ΔRING) to examine the necessity of the RING domain and found that *Trim41*^ΔRING/ΔRING^ phenocopied *Trim41^neo/neo^* (i.e., KO). Surprisingly, the ΔRING-TRIM41 accumulated on the SYCP3-overloaded regions. The ΔRING-TRIM41 accumulation also correlated with the chaotic X chromosome synapsis.

The most striking phenotype of *Trim41* KO spermatocytes was SYCP3 overloading. In addition, some X chromosomes exhibited multilayered SYCP3 signals, clearly demonstrating self-synapsis of the X chromosome. Although we could not specify, this self-synapsis might be inter-sister synaptonemal complex assembly like those found in REC8 KO spermatocytes (Ishiguro et al., 2014; Ishiguro and Watanabe, 2016; Xu et al., 2005). However, given that we also observed multilayer SYCP1 signals in the X chromosome, we should also consider the possibility of extensive heterologous self-synapsis like those seen in domestic dog pachytene spermatocytes (Federici et al., 2015). The latter possibility is corroborated by another outstanding phenotype of the non-homologous synapsis between the X chromosome and autosomes. Further studies are needed to determine the relationship between these two phenotypes (i.e., SYCP3 overloading and abnormal synapsis configuration; causal relationship or simultaneous phenomena) and whether these phenotypes are direct influences of TRIM41 deficiency or compensatory changes.

To examine which chromosomes exhibited SYCP3 overloading in *Trim41* KO spermatocytes, we performed multicolor FISH combined with SYCP3 immunostaining and found that the X chromosome was mainly affected. On the other hand, autosomes saw fewer SYCP3 overloading and no biases to specific chromosomes. A possible reason for the X chromosome bias is the unsynapsed nature during the pachytene stage. This inference agrees that only a tiny population of germ cells underwent SYCP3 overloading in *Trim41* KO females, whose X chromosomes are fully synapsed during prophase I progression. Therefore, *Trim41*-deficient XO meiosis might be an intriguing future consideration.

To examine the molecular function of TRIM41, we also produced *Trim41^ΔRING/ΔRING^* mice by replacing 2–61 aa residues with HA-tag sequence, and we found that the HA immunostaining signals on overloaded-SYCP3 region. As the HA signals appeared only when the cells had no functional TRIM41 (i.e. *Trim41^ΔRING/ΔRING^* or *Trim41^neo/ΔRING^*), we concluded that ΔRING-TRIM41 accumulated on the SYCP3-overloaded regions. This observation suggested that ΔRING-TRIM41 kept targeting the unubiquitinated substrates, and therefore TRIM41 exerts its ubiquitin ligase E3 activity on the chromosome axes. However, we could not narrow down the substrate candidate by IP-Mass analysis. Also, we did not see the interaction between TRIM41 and SYCP3. As the weak/transient binding between enzymes and substrates tends to be challenging to capture in IP analysis (Trinkle-Mulcahy, 2019), the recently developed proximity-dependent biotin identification (BioID) technique (Kim et al., 2016; Roux et al., 2012) could be a future consideration in substrate identification.

In summary, our result showed that *Trim41* is essential for preventing SYCP3 overloading and X chromosome chaotic synapsis. Further studies for substrate identification will unveil the exact molecular functions of TRIM41 and the mechanism for safeguarding X chromosome chaotic synapsis in male meiosis.

## Material & methods

### Animals

All animal experiments were approved by the Animal Care and Use Committee of the Research Institute for Microbial Diseases, Osaka University (#Biken-AP-R03-01-0). Animals were housed in a temperature-controlled environment with 12 h light cycles and free access to food and water. B6D2F1 (C57BL/6 × DBA2; Japan SLC, Shizuoka, Japan) mice and ICR (SLC) were used as embryo donors; B6D2F1 were used for mating and wild-type control; C57BL6/N (SLC) mice were used to collect RNA for cloning.

### Generation of Trim41 KO mice

*Trim41* KO mice were generated by transfecting EGR-G101 (Fujihara et al., 2013) with a targeting vector. Then, potentially targeted ES cell clones were selected with neomycin. Correctly targeted ES cell clones and germ-line transmission were confirmed via PCR using primers (GeneDesign, Osaka, Japan) listed in Table S1. Finally, the heterozygous KO mice were mated with B6D2F1 and then maintained by sibling mating. The B6D2 KO mouse line is available from the Riken BioResource Center (Riken BRC, Tsukuba, Japan; #11261) and the Center for Animal Resources and Development, Kumamoto University (CARD, Kumamoto, Japan; #3019).

### Generation of Trim41 transgenic mice

The mouse *Trim41* cDNA (ENSMUST00000047145) tagged with PA and 1D4 was inserted under the control of the mouse Clgn promoter (Addgene #173686) (Ikawa et al., 2001; Watanabe et al., 1995). The *Trim41* cDNA inserted plasmids are deposited at Riken BRC and Addgene. After linearization, the DNA construct (2.16 ng/μL; 0.54 ng/μL/kbp) was injected into the pronucleus of fertilized eggs. The injected eggs were transplanted into the oviduct ampulla of pseudopregnant mice (ICR; 10 embryos per ampulla). After 19 days, pups were delivered through Caesarean section and placed with foster mothers (ICR). For the rescue experiment, F0 Tg mice were mated with *Trim41* KO mouse line. The mouse colony with the transgene and KO allele was maintained by sibling mating. The genotyping primers (GeneDesign) are listed in Table S1. The transgenic mouse line is available from Riken BRC (#11216) and CARD (#3019).

### Generation of ΔRING-Trim41 mice

*ΔRING-Trim41* mice were generated by microinjection described previously (Oura et al., 2020). First, a gRNA solution was prepared by annealing two tracrRNAs (Sigma-Aldrich, St. Louis, MO, USA) and crRNA (Sigma-Aldrich). The target genomic sequences are listed in Table S1. Then, the gRNA solution and Cas9 nuclease solution (Thermo Fisher Scientific, Waltham, MA, USA) were mixed: 40 ng/μL gRNA each and 108 ng/μL Cas9 nucleases. The obtained complex was then microinjected into fertilized eggs (B6D2F1) using a programmable microinjector (FemtoJet 4i, Eppendorf, Hamburg, Germany). The microinjected eggs were then transplanted into the oviduct ampulla of pseudopregnant mice (ICR) on the following day. After 19 days, pups were delivered through Caesarean section and placed with foster mothers (ICR). To generate heterozygous mutant mice, F0 mice were mated with WT B6D2F1. Mouse colonies with the desired mutation were maintained by sibling mating. The genotyping primers (GeneDesign) are listed in Table S1. The mutant mouse line is available from Riken BRC (#11041) and CARD (#2948).

### Bacterial strains

*Escherichia coli* (*E. coli*) strain DH5α (Toyobo, Osaka, Japan) and BL21(de3) pLysS (C606003, ThermoFisher Scientific) were used for DNA cloning and protein expression, respectively. E. coli cells were grown in LB or 2×YT medium containing 100 mg/L ampicillin and were transformed or cloned using standard methods.

### Production of anti-TRIM41 antibody

Monoclonal antibody against TRIM41 was produced as previously described (Oura et al., 2021). The DNA encoding mouse TRIM41 (residue 35-85 aa, NP_663352.2) was inserted into pGEX6p-1 (GE healthcare), and the expression vector was transformed into *E. coli* strain BL21 (de3) pLysS (C606003, Thermo Fisher Scientific). GST-TRIM41 was purified using Glutathione Sepharose 4B (GE Healthcare). The GST tag was removal by PreScission protease (27084301, GE Healthcare) and Glutathione Sepharose 4B affinity subtraction purification. The purified TRIM41 protein with a complete adjuvant was injected into female rats. After 17 days of injection, lymphocytes were collected from iliac lymph nodes for hybridomas generation (Kishiro et al., 1995; Sado et al., 2006). The hybridomas were cloned by a limited dilution and screened by ELISA against recombinant TRIM41 and immunoblotting against testis lysate. A monoclonal antibody from hybridoma clone #33-16 was used in this study. As a side note, the #33-16 antibody also recognized ΔRING-TRIM41 missing 2-61 aa residues, showing that the epitope of #33-16 is in 59-85 aa residues (two or three mismatches might be acceptable).

### Genotype analysis

PCR was performed using KOD FX neo (KFX-201, TOYOBO). The primers (GeneDesign) for each gene are summarized in Table S1. PCR products were purified using a Wizard SV Gel and PCR Clean-Up System (Promega, Madison, WI, USA) kit for Sanger sequencing.

### Sequence comparison analysis

Amino acid sequences of TRIM41 were obtained from the NCBI Entrez Protein database. Clustal W2.1 was used for multiple sequence alignment (Larkin et al., 2007).

### Immunoblotting

Proteins from testis were extracted using NP40 lysis buffer [50mM Tris-HCl (pH 7.5), 150 mM NaCl, 0.5% NP-40, 10% Glycerol]. Proteins were separated by SDS-PAGE under reducing conditions and transferred to polyvinylidene fluoride (PVDF) membrane using the Trans Blot Turbo system (BioRad, Munich, Germany). After blocking with 10% skim milk (232100, Becton Dickinson, Cockeysville, MD, USA), the membrane was incubated with primary antibody overnight at 4°C, and then incubated with HRP-conjugated secondary antibody for 1 h at room temperature. Chemiluminescence was detected by ECL Prime Western Blotting Detection Reagents (RPN2232, GE Healthcare, Chicago, IL, USA) using the Image Quant LAS 4000 mini (GE Healthcare). The antibodies used in this study are listed in Table S2.

### Fertility analysis

To examine fertility, sexually mature male mice were caged with wild-type females (B6DF1) for at least three months. The vaginal plugs and pup’s numbers were recorded at approximately 10 AM to determine the number of copulations and litter size. Numerical data is available in Table S3.

#### Morphological and histological analysis of testis and epididymis

To observe gross testis morphology and measure testicular weight, mice over 11-week-old age were euthanized after measuring their body weight. Numerical data is available in Table S3. The whole testis was observed using BX50 and SZX7 (Olympus, Tokyo, Japan) microscopes. For histological analysis, testes were fixed in Bouin’s fixative solution (16045-1, Polysciences, Warrington, PA, USA) at 4°C O/N, dehydrated in increasing ethanol concentrations and 100% xylene, embedded in paraffin, and sectioned (5 μm). The paraffin sections were hydrated with 100% Xylene and decreasing ethanol concentrations, treated with 1% periodic acid (26605-32, Nacalai Tesque, Kyoto, Japan) for 10 min, incubated with Schiff’s reagent (193-08445, Wako) for 20 min, counterstained with Mayer’s hematoxylin solution (131-09665, Wako) for 3 min, dehydrated in increasing ethanol concentrations, and finally mounted with Permount (SP15-100-1, Ferma, Tokyo, Japan). The sections were observed using a BX53 (Olympus) microscope. Seminiferous tubule stages were identified based on the morphological characteristics of the germ cell nuclei (Ahmed and de Rooij, 2009).

### Apoptosis detection in testicular section

TdT-mediated dUTP nick end labeling (TUNEL) staining was carried out with In Situ Apoptosis Detection Kit (MK500, Takara Bio Inc., Shiga, Japan), according to the manufacturer’s instruction. Briefly, testes were fixed with Bouin’s fixative, embedded in paraffin, and sectioned (5 μm). After paraffin removal, the slides were boiled in citrate buffer (pH 6.0; 1:100; ab93678, abcam, Cambridge, UK) for 10 min and incubated in 3% H_2_O_2_ at room temperature for 5 min for endogenous peroxidase inactivation, followed by a labeling reaction with TdT enzyme and FITC-conjugated dUTP at 37°C for 1 h.

For chromogenic detection of apoptosis, the sections were incubated with HRP-conjugated anti-FITC antibody at 37°C for 30 min. The section was then incubated in ImmPACT DAB (SK-4105, Vector Laboratories, Burlingame, CA, USA) working solution, counterstained with Mayer’s hematoxylin solution for 3 min, dehydrated in increasing ethanol concentrations, and finally mounted with Permount. The sections were observed using a BX53 (Olympus) microscope. Seminiferous tubule stages were identified based on the morphological characteristics of the germ cell nuclei (Ahmed and de Rooij, 2009). Numerical data is available in Table S3.

### Immunostaining of surface chromosome spread

Spread nuclei from spermatocytes were prepared as previously described (Oji et al., 2020; Oura et al., 2021). Seminiferous tubules were unraveled using forceps in ice-cold DMEM (11995065, Thermo Fisher Scientific) and incubated in 1 mg/mL collagenase (C5138, Sigma-Aldrich) in DMEM (20 mL) at 37°C for 15 min. After three washes with DMEM, the tubules were transferred to 20 mL trypsin/DNaseI medium [0.025 w/v% trypsin, 0.01 w/v% EDTA, 10U DNase in DMEM] and incubated at 37 °C for 10 min. After adding 5 mL of heat-inactivated FCS and pipetting, the solution was filtered through a mesh (59 μm; N-N0270T, NBC Meshtec inc., Tokyo, Japan) to remove tubule debris. The collected testicular cells were resuspended in hypotonic solution [100 mM sucrose] and 10μL of the suspension was dropped onto a slide glass with 100 μL of fixative solution [1% PFA, 0.1% (v/v) Triton X-100]. The slides were then air-dried and washed with PBS containing 0.4% Photo-Flo 200 (1464510, Kodak Alaris, NY, USA) or frozen for longer storage at −80°C.

The spread samples were blocked with 10% goat serum in PBS and then incubated with primary antibodies overnight at 4°C in blocking solution. After incubation with AlexaFlour 488/546-conjugated secondary antibody (1:200) at room temperature for 1 h, samples are counterstained with Hoechst 33342 and mounted with Immu-Mount. The samples were observed using a BX53 (Olympus) microscope. The antibodies used in this study are listed in Table S2. Numerical data is available in Table S3.

#### Giemsa staining of metaphase I chromosome spreads

For preparing metaphase chromosome spreads, seminiferous tubules were unraveled using forceps in ice-cold PBS and transferred to a 1.5-mL tube with 1 mL of accutase (12679–54, Nacalai Tesque), followed by clipping the tubules, and a 5 min incubation at room temperature. After filtration with a mesh (59 μm; N-N0270T, NBC Meshtec inc.) and centrifugation, the cells were resuspended in 8 mL of hypotonic solution [1% sodium citrate] and incubated for 5 min at room temperature. The suspension was centrifuged and 7 mL of supernatant was aspirated. The cells were then resuspended in the remaining 1 mL of supernatant and 7 mL of Carnoy’s Fixative [75% Methanol, 25% Acetic Acid] were added gradually while shaking. After 2 washes with Carnoy’s Fixative, the cells were resuspended in ~ 0.5 mL of Carnoy’s Fixative and dropped onto a wet glass slide. The slide was stained with Giemsa Stain Solution (079–04391, wako) and observed using a BX53 (Olympus) microscope.

### Multicolor fluorescence in situ hybridization (mFISH) followed by SYCP3 immunostaining

Testicular germ cells were suspended in 3 mL of hypotonic solution [0.075% potassium chrolide] and incubated for 20 min at room temperature. Then the cells were mixed with 1 mL of Carnoy’s Fixative [75% Methanol, 25% Acetic Acid] to the suspension for fixing. After 5 mL of Carnoy’s Fixative addition, the suspension was again mixed and centrifuged at 1,500 rpm for 5 minutes. After three washes with 5 mL of Carnoy’s Fixative, the fixed cells were subjected to metaphase spreader Hanabi (ADSTEC, Funabashi, Japan) for spread sample preparation.

For multicolor FISH analysis, the spread samples were hybridized with 21XMouse (MetaSystems, Altlussheim, Germany) according to the manufacturer’s protocol. For denaturation of the nuclear DNA, the spread samples were incubated in 2×SSC for 30 min at 70°C and then treated with 0.07M NaOH for 1 min at room temperature. The denatured samples were washed with 0.1×SSC and 2×SSC for 1 min at 4°C, respectively, and then dehydrated with 70%, 95%, and 100% ethanol. Multicolor FISH probes were denatured for 5 min at 75°C and applied to the spread samples. After hybridization for 48 h at 37°C in a humidified chamber, the spread samples were treated with 0.4× SSC for 2 min at 72°C, washed in 2×SSC containing 0.05% Tween20 for 30 sec at room temperature, and rinsed with distilled water.

Immunostaining of the spread nuclei with SYCP3 was performed described above. After incubation of the spread samples with primary and secondary antibodies, the slides were covered by a coverslip with DAPI/Antifade (MetaSystemes, Altlussheim, Germany) and observed under a fluorescent microscope. Fluorescence images were captured using a high-sensitive digital camera (α7s, SONY, Tokyo, Japan), and the chromosome numbering of each synaptonemal complex was determined based on the fluorescence color.

### Immunoprecipitation

Proteins from seminiferous tubules were extracted using NP40 lysis buffer [50 mM Tris-HCl (pH7.5), 150 mM NaCl, 0.5% NP-40, 10% Glycerol]. Protein lysates were mixed with 20 μL Protein G-conjugated magnetic beads (DB10009, Thermo Fisher Scientific) with 2.0 μg antibody. The immune complexes were incubated for 1 h at 4°C and washed 3 times with NP40 lysis buffer. Co-immunoprecipitated products were then eluted by resuspension in 2x SDS sample buffer [125 mM Tris-HCl (pH6.8), 10% 2-mercaptoethanol, 4% sodium dodecyl sulfate (SDS), 10% sucrose, 0.01% bromophenol blue] and 10 min incubation at 70°C. The antibodies used in this study are listed in Table S2.

### Mass spectrometry and data analysis

Before MS analysis, half of the eluted amount was subjected to SDS-PAGE and silver staining (06865-81, Nacalai Tesque). The remaining half amount was subjected to mass spectrometry (MS) analysis. The proteins were reduced with 10 mM dithiothreitol (DTT), followed by alkylation with 55 mM iodoacetamide, and digested by treatment with trypsin and purified with a C18 tip (GL-Science, Tokyo, Japan). The resultant peptides were subjected to nanocapillary reversed-phase LC-MS/MS analysis using a C18 column (25 cm × 75 um, 1.6 μm; IonOpticks, Victoria, Australia) on a nanoLC system (Bruker Daltoniks, Bremen, Germany) connected to a tims TOF Pro mass spectrometer (Bruker Daltoniks) and a modified nano-electrospray ion source (CaptiveSpray; Bruker Daltoniks). The mobile phase consisted of water containing 0.1% formic acid (solvent A) and acetonitrile containing 0.1% formic acid (solvent B). Linear gradient elution was carried out from 2% to 35% solvent B for 18 min at a flow rate of 400 nL/min. The ion spray voltage was set at 1.6 kV in the positive ion mode. Ions were collected in the trapped ion mobility spectrometry (TIMS) device over 100 ms and MS and MS/MS data were acquired over an *m/z* range of 100-1,700. During the collection of MS/MS data, the TIMS cycle was adjusted to 1.1 s and included 1 MS plus 10 parallel accumulation serial fragmentation (PASEF)-MS/MS scans, each containing on average 12 MS/MS spectra (>100 Hz), and nitrogen gas was used as the collision gas.

The resulting data were processed using DataAnalysis version 5.1 (Bruker Daltoniks), and proteins were identified using MASCOT version 2.6.2 (Matrix Science, London, UK) against the SwissProt database. Quantitative value (available in Table S4) and fold exchange were calculated by Scaffold4 (Proteome Software, Portland, OR, USA) for MS/MS-based proteomic studies.

### Statistics and Reproducibility

All error bars indicated standard deviation. The sample numbers were described in each legend or/and in the figure panel.

## Acknowledgments

We wish to thank the members of both the Department of Experimental Genome Research, Animal Resource Center for Infectious Diseases, and NPO for Biotechnology Research and Development for experimental assistance. We also thank Dr. Kusakabe and Dr. Tateno at Department of Biological Sciences, Asahikawa Medical University for technical consultation of mFISH experiments.

## Data availability

The authors declare that the data supporting this study’s findings are available from the corresponding author upon request. Numerical data that underlies graphs were listed in Table S3.

## Supplemental information

**Fig. S1: *Trim41* is an evolutionally conserved gene expressed highly in pachytene germ cells.**

(A) The percentage of TF342569 annotated species based on TreeFam database (Release 9; http://www.treefam.org/). Dark and light green show species with and without TF342569 annotation, respectively. (B) Protein sequence comparison of TRIM41 in big brown bat (XP_027991236.1), cat (XP_003980687.1), cattle (NP_001193094.1), chimpanzee (XP_016809993.1), dog (XP_038536985.1), human (NP_291027.3), mouse (NP_663352.2), pig (XP_020939058.1), and rat (NP_001128209.1). (C) The *Trim41* expression profile between testicular cells based on published single-cell RNA sequencing data, visualized by 10 x genomics Loupe Browser. UMI means Unique Molecular Identifiers. (D) The *Trim41* expression profile between embryonic female germ cells based on published single-cell RNA sequencing data.

**Fig. S2: Expression confirmation of *Clgn-Trim41 Tg***

(A) RT-PCR using postnatal testis of *Trim41* KO-Tg male mice. (B) Immunoblotting analysis with an anti-PA antibody. RT means reverse transcription. The black arrow indicates a specific band in Tg. (C) Immunoblotting analysis with an anti-TRIM41 antibody raised against a recombinant TRIM41 protein (35-85 amino acid residues). The black and red arrows indicate a specific band in Tg and WT, respectively.

**Fig. S3: Immunostaining of zygotene–pachytene spermatocytes, related to Fig 3A-C.**

(A and B) Additional immunostaining images of zygotene–pachytene spermatocytes from Het (A) and KO (B) male mice. The white dashed line shows nuclei of different cells. The white arrowhead indicates SYCP3 overloading. The images were arranged in the order of meiotic progression from the zygotene stage (top) to the early pachytene stage (bottom).

**Fig. S4: Immunostaining of meiotic cohesins.**

(A and B) SYCP3/REC8 immunostaining of spread nuclei from prophase spermatocytes collected from adult WT (A) and KO (B) male mice. (C and D) SYCP3/RAD21L immunostaining of spread nuclei from prophase spermatocytes collected from adult WT (A) and KO (B) male mice.

**Fig. S5: Phenotypical analysis of *Trim41* KO females**

(A) Gross morphology of female gonads from E15.5 fetus. (B) Immunostaining of female gonad sections. White arrows indicate intense SYCP3 signals. (C) Ovarian histology of *Trim41* Het and KO females. The ovaries were collected 10 h after intraperitoneal administration of hCG.

**Fig. S6: Immunostaining of prophase I spermatocytes, related to Fig 6A and B.**

(A and B) Immunostaining of spread nuclei from prophase spermatocytes collected from adult ΔR-Het (A) and ΔR-Homo (B) male mice. At least three male mice were analyzed. (C) The percentage of each meiotic prophase stage in immunostained spread nuclei samples in A and B. The numerical data is available in Table S3.

**Fig. S7: Additional images for ΔRING-TRIM41 accumulation, related to Fig 6C-F.**

(A) SYCP3/HA immunostaining of zygotene–diplotene spermatocytes from) ΔR-Homo male mice. White and blue arrowheads indicate HA signals on and outside of SYCP3 axes, respectively. At least three male mice were analyzed. (B–E) SYCP3/TRIM41 immunostaining of pachytene spermatocytes from Het (B), KO (C), ΔR-Het (D), and ΔR-Homo (E) male mice. White arrowheads indicate HA signals on SYCP3 axes. At least three male mice were analyzed.

**Fig. S8: Additional images for SYCP1 and BRCA1 immunostaining, related to Fig 6G and H.**

(A) SYCP1/HA immunostaining of surface chromosome spread from ΔR-Homo testis. Magenta and yellow arrowheads indicate HA signals on the SYCP1 positive/negative boundary and the non-homologous axis interface, respectively. At least three male mice were analyzed. (B) BRCA1/HA immunostaining of surface chromosome spread from ΔR-Homo testis. Red arrowheads indicate HA signals on the BRCA1 negative part of the X chromosome axes. At least three male mice were analyzed.

**Fig. S9: Immunoprecipitation for substrate identification.**

(A and B) Immunoprecipitation using an anti-TRIM41 antibody and WT testis lysate. TRIM41 immunoblotting (A) validated the immunoprecipitation capability of the anti-TRIM41 antibody and the success of immunoprecipitation experiments. Red and green arrowheads indicate TRIM41 and SYCP3, respectively. (C and D) Immunoprecipitation using an anti-TRIM41 antibody and ΔR-Homo testis lysate. HA immunoblotting (C) validated the success of immunoprecipitation experiments. Black and green arrowheads indicate HA-ΔRING-TRIM41 and SYCP3, respectively. (E and F) Immunoprecipitation using an anti-HA antibody and ΔR-Homo testis lysate. HA immunoblotting (E) validated the success of immunoprecipitation experiments. Black and green arrowheads indicate HA-ΔRING-TRIM41 and SYCP3, respectively. (G and H) Mass analysis of co-IPed elutes using anti-TRIM41 (G) and anti-HA (H) antibodies. Proteins with 0 spectra in KO lysate were extracted and summarized in the table. The whole proteomics data are available in Table S4. (I) Proteins detected more than twice in G and H were summarized.

**Table. S1: Primers and gRNAs used in this study.**

**Table. S2: Antibodies used in this study.**

**Table. S3: Numerical data that underlies graphs.**

**Table. S4: The quantitative value of mass spectrometry analysis.**

