## supplemental figures for "*Trim41* is essential for preventing X chromosome chaotic synapsis in male mice"

### Supplementary Figure1

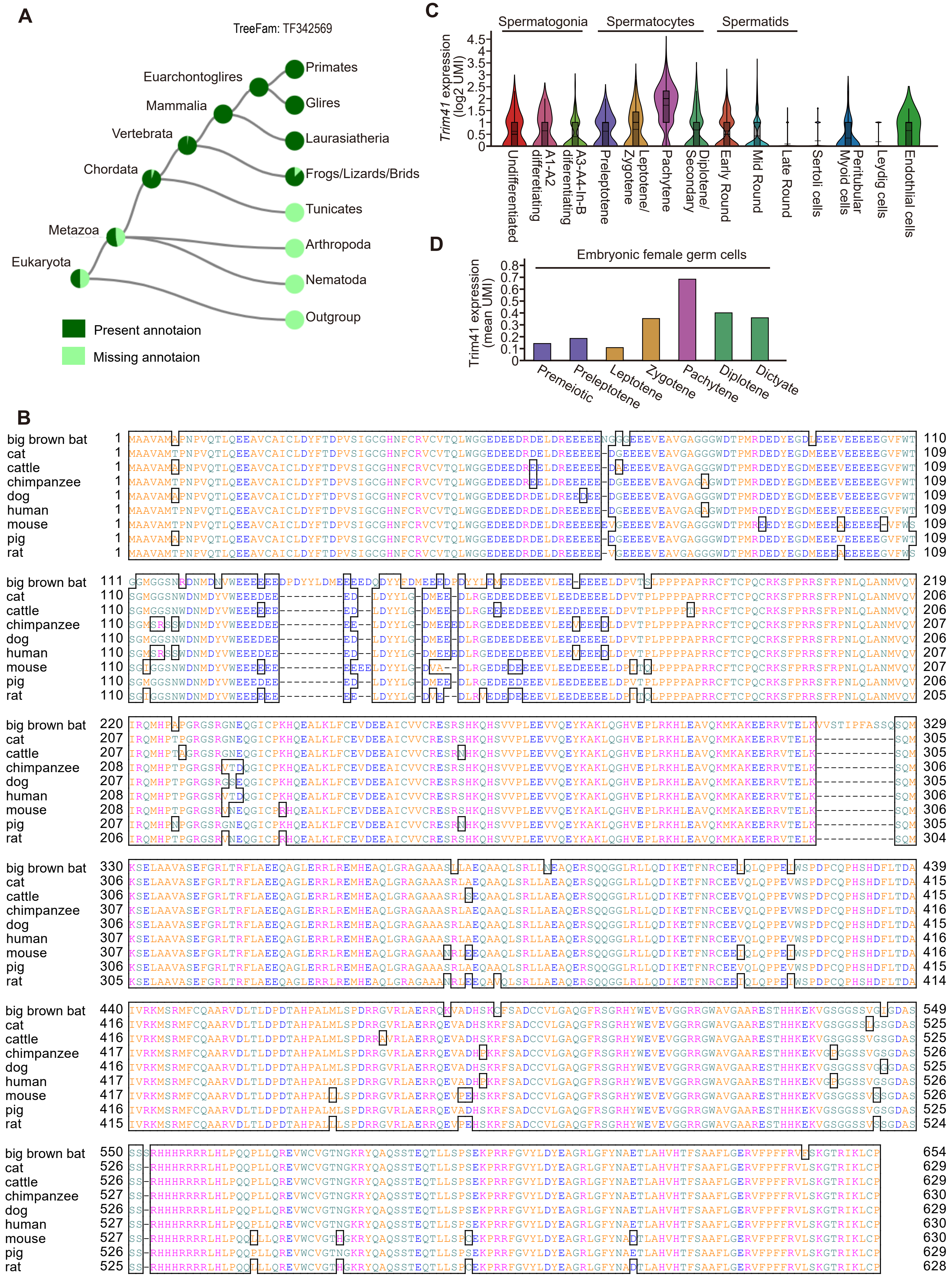

### Supplementary Figure 2

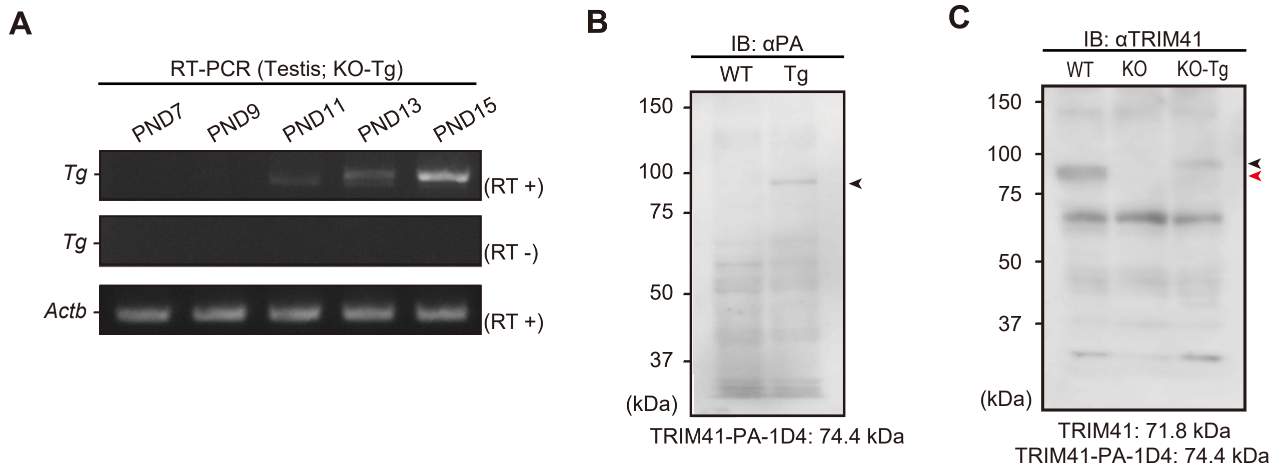

### Supplementary Figure 3

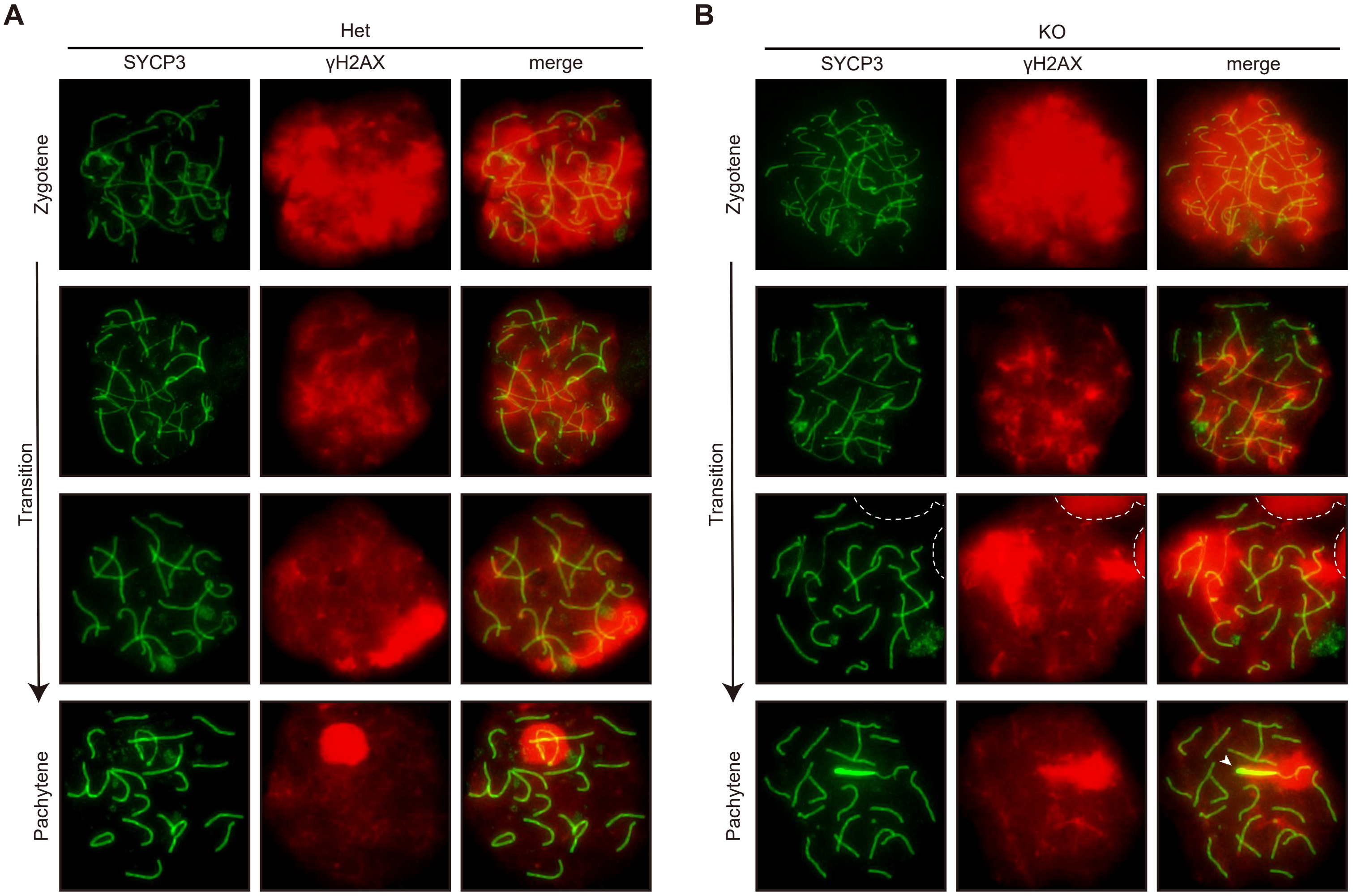

### Supplementary Figure 4

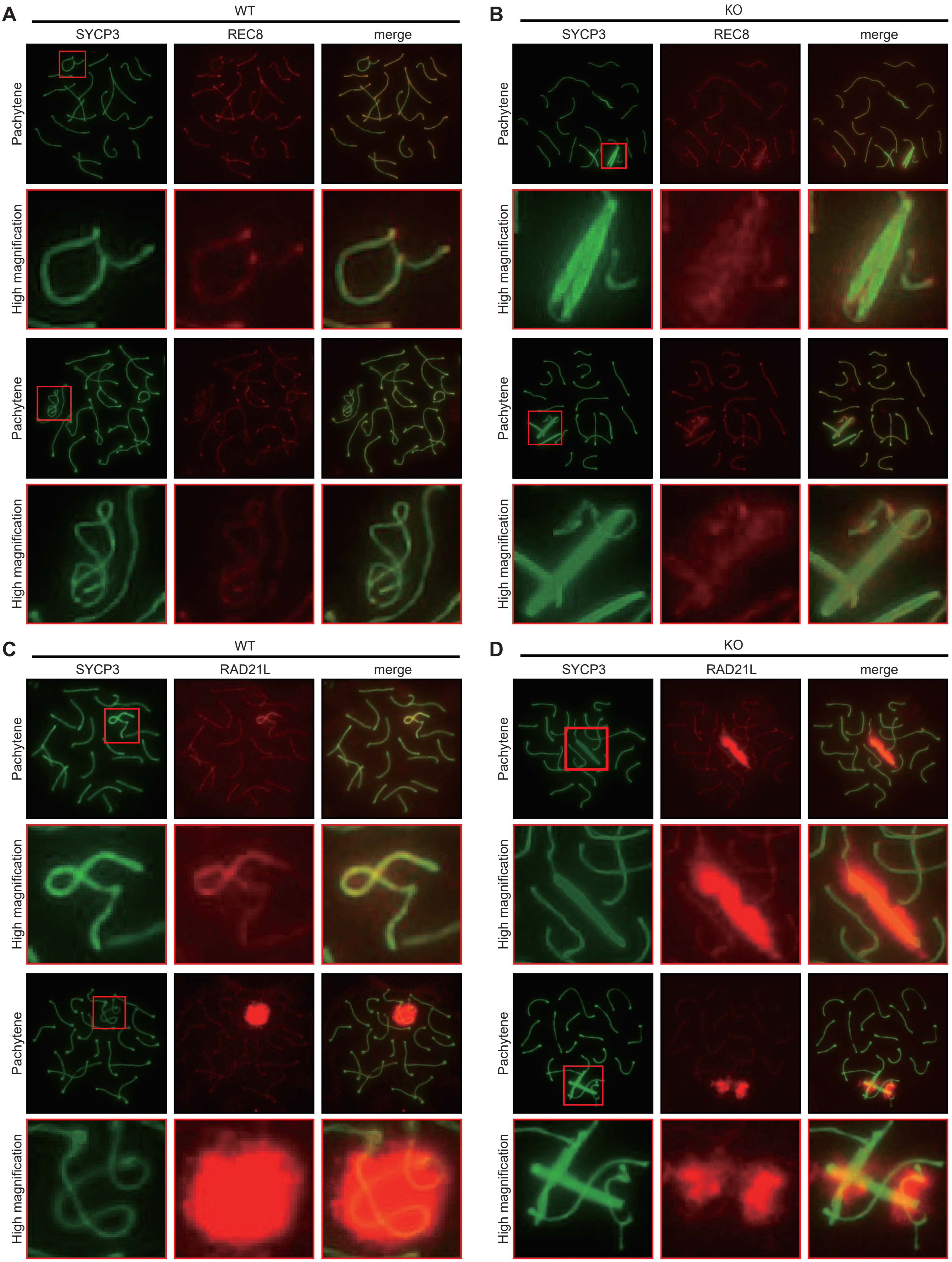

### Supplementary Figure 5

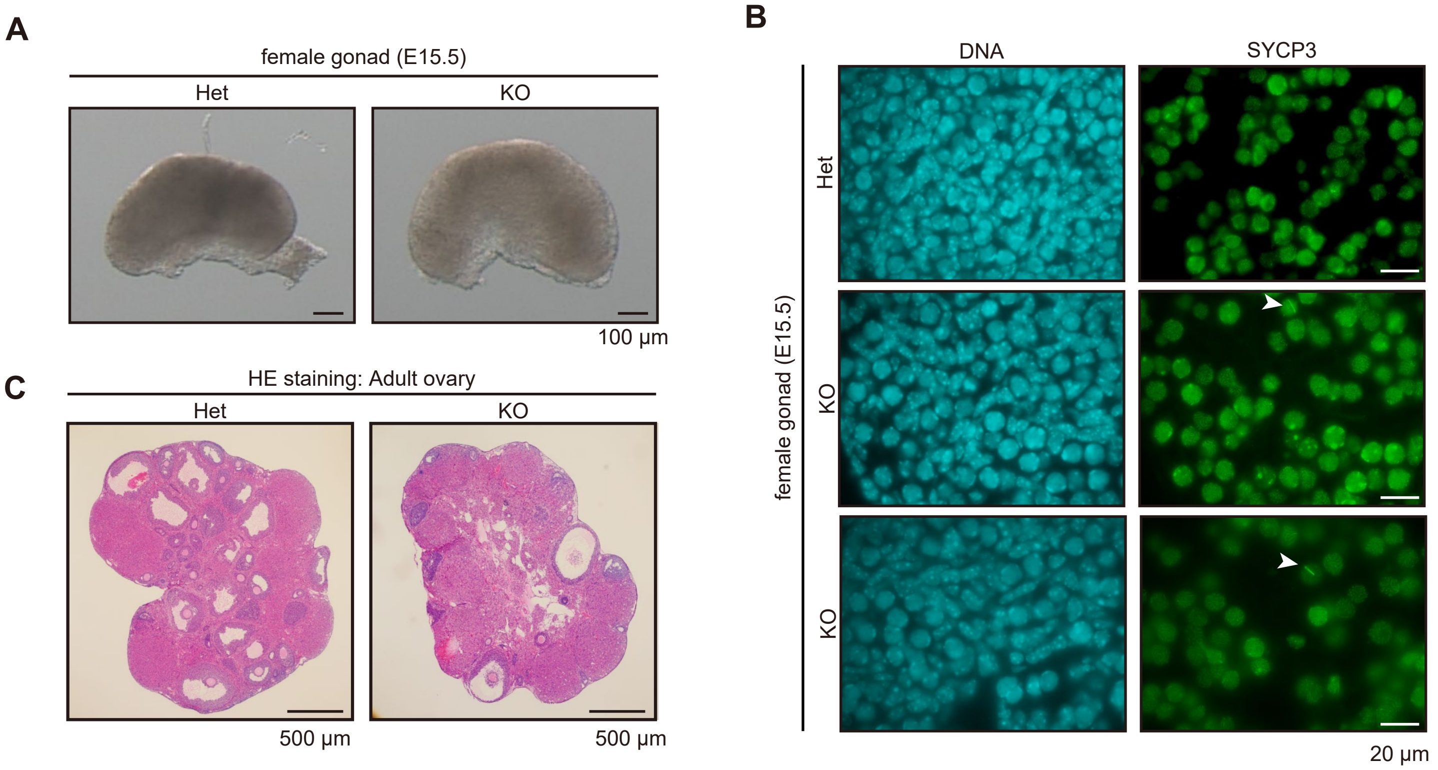

### Supplementary Figure 6

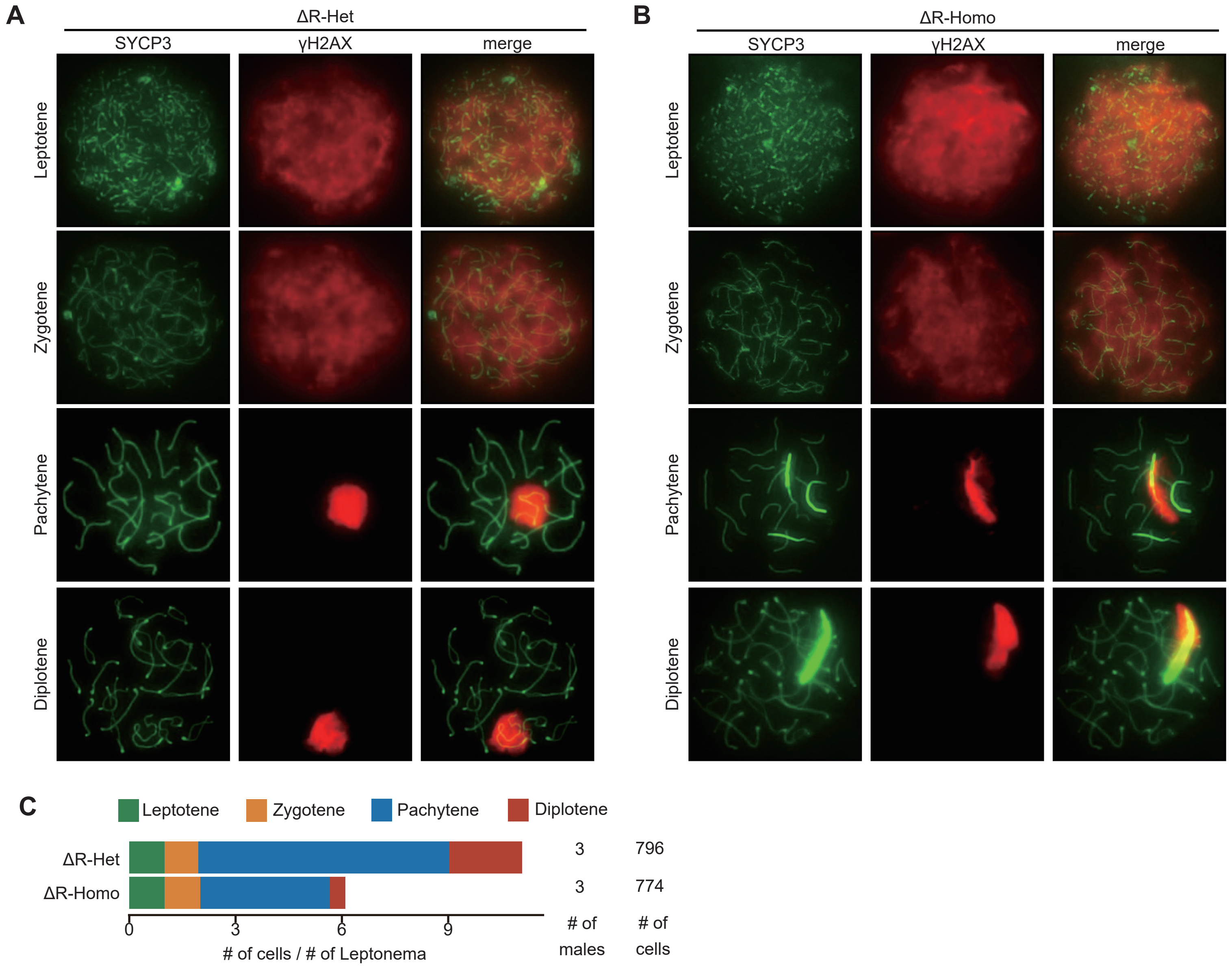

### Supplementary Figure 7

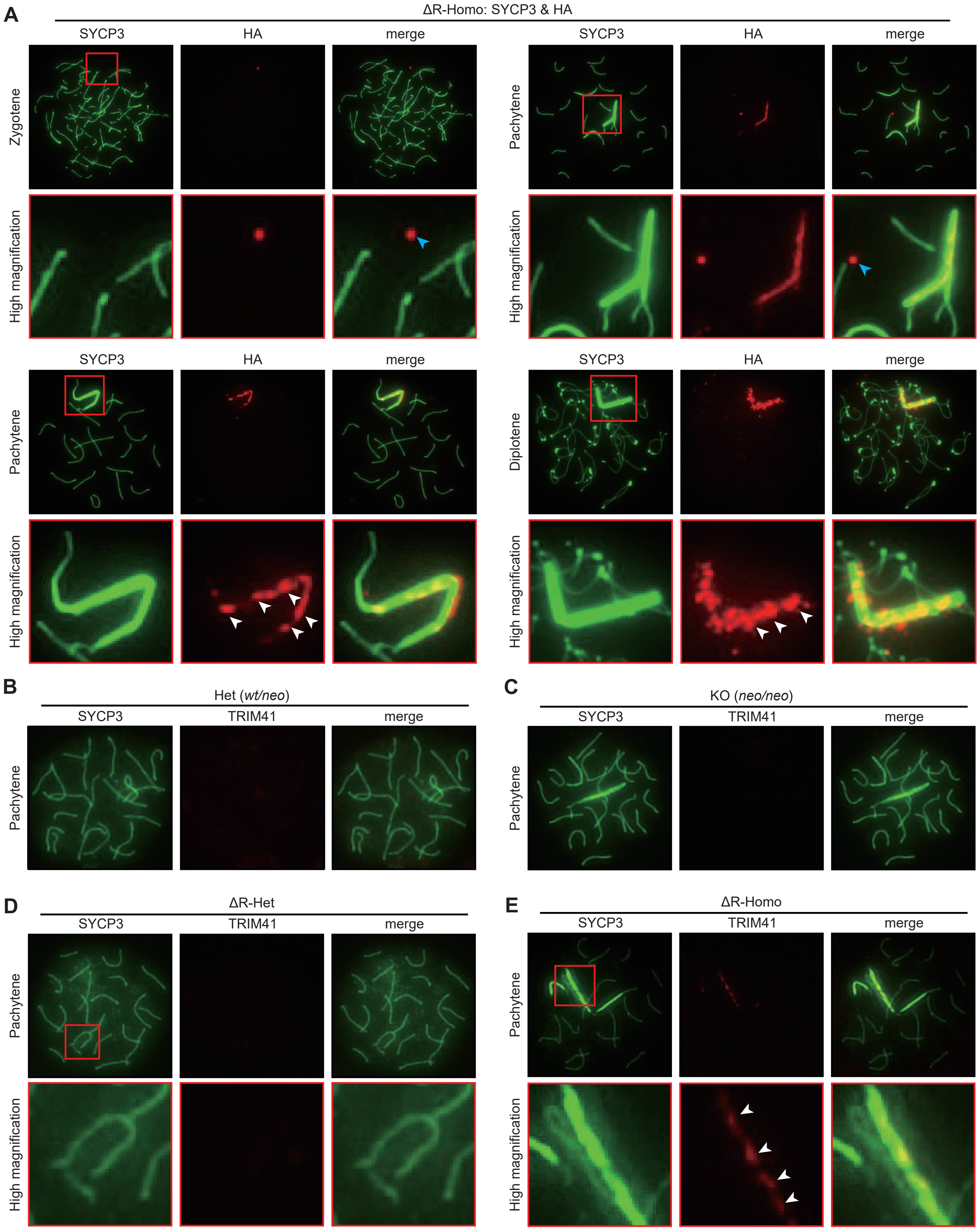

### Supplementary Figure 8

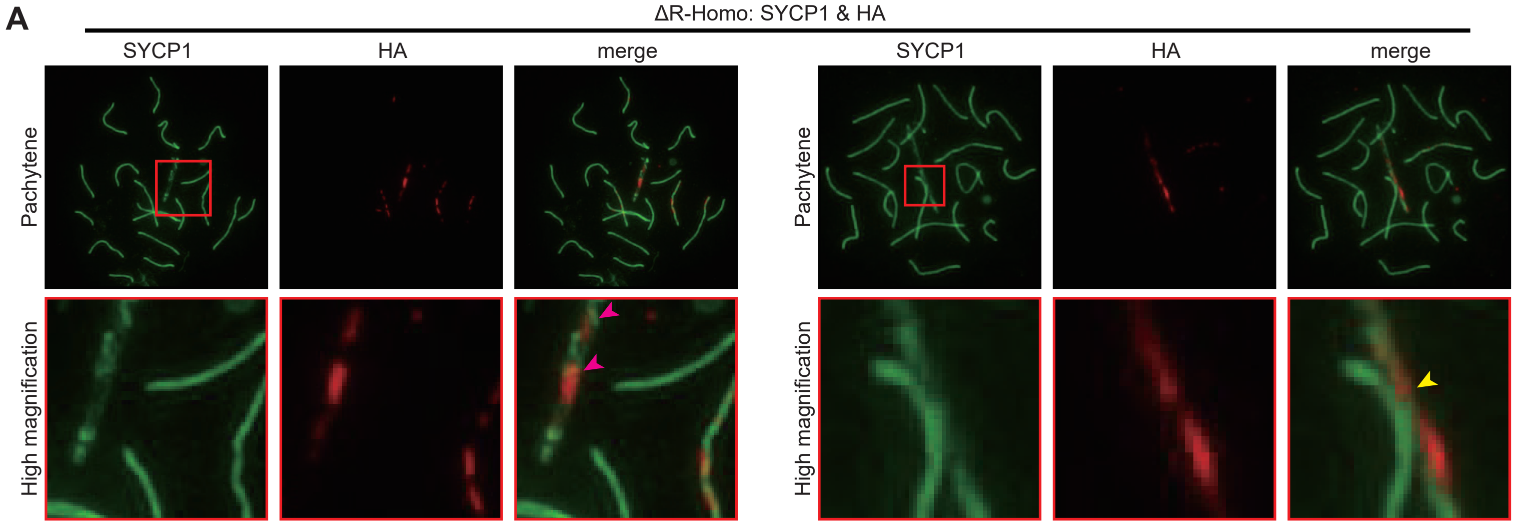


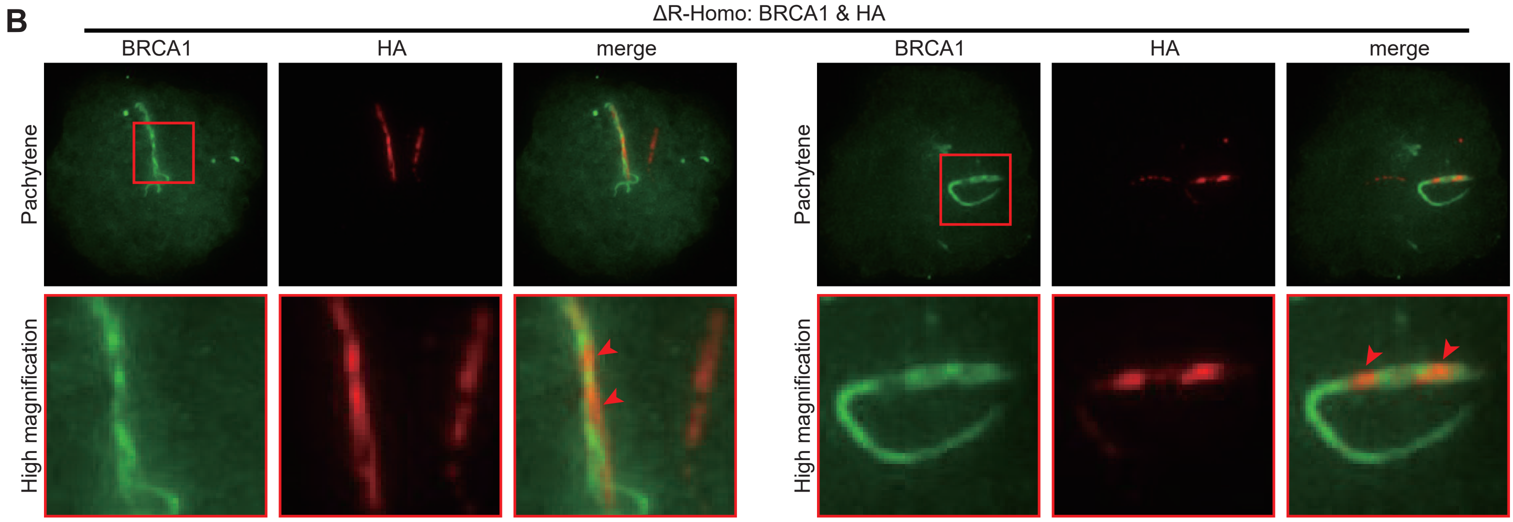

### Supplementary Figure 9

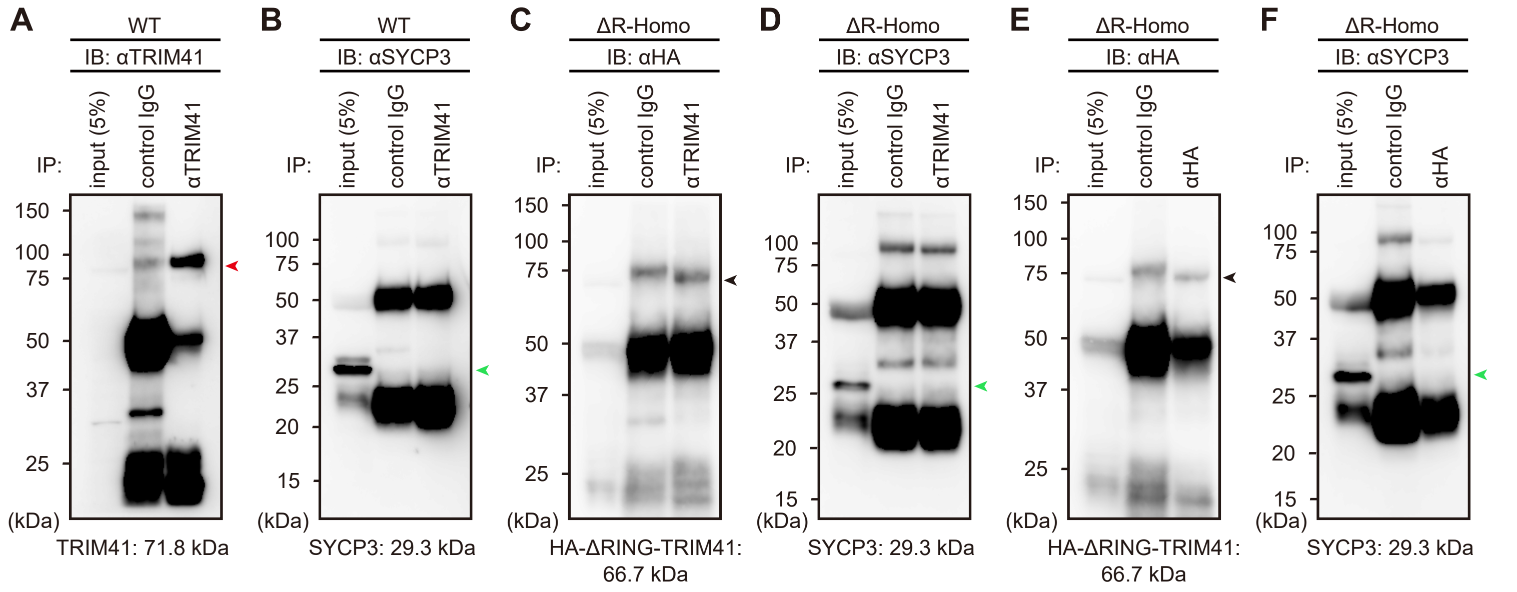


**G**

| Protein | MW | Accession Number | Quantitative Value (Normalized Total Spectra) IP: αTRIM41 WT | ΔR-Homo | KO |
|---|---|---|---|---|---|
| TRIM41 | 72 kDa | Q5NCC3 | 22.724 | 1.1042 | 0 |
| CCDC40 | 137 kDa | Q8BI79 | 6.3627 | 1.1042 | 0 |
| CCDC42 | 38 kDa | Q5SV66 | 6.3627 | 0 | 0 |
| NT5C1B | 65 kDa | Q91YE9 | 6.3627 | 0 | 0 |
| | 19 kDa | Q5BN45 | 5.4538 | 0 | 0 |
| CFAP91 | 92 kDa | Q8BRC6 | 5.4538 | 0 | 0 |
| DNAH12 | 356 kDa | Q3V0Q1 | 5.4538 | 0 | 0 |
| CCDC39 | 110 kDa | Q9D5Y1 | 5.4538 | 0 | 0 |
| DRC1 | 89 kDa | Q3USS3 | 5.4538 | 0 | 0 |
| MAP7D3 | 98 kDa | A2AEY4 | 4.5448 | 0 | 0 |
| TSSK4 | 37 kDa | Q9D411 | 4.5448 | 0 | 0 |
| PRKAR2A | 45 kDa | P12367 | 4.5448 | 0 | 0 |
| DRC7 | 103 kDa | Q6V3W6 | 4.5448 | 0 | 0 |
| | 35 kDa | Q3U1D9 | 4.5448 | 0 | 0 |
| ODAD1 | ? | Q3UX62 | 3.6358 | 0 | 0 |
| | 21 kDa | Q9DAR0 | 3.6358 | 0 | 0 |
| SPICE1 | 96 kDa | Q8C804 | 3.6358 | 0 | 0 |
| SPATA6 | 56 kDa | Q3U6K5 | 3.6358 | 0 | 0 |
| DYNLRB2 | 11 kDa | Q9DAJ5 | 3.6358 | 0 | 0 |
| STK36 | 144 kDa | Q69ZM6 | 3.6358 | 0 | 0 |
| CCIN | 67 kDa | Q8CDE2 | 2.7269 | 0 | 0 |
| HSP90AB1 | 83 kDa | P11499 | 2.7269 | 1.1042 | 0 |
| | 68 kDa | Q9D5Y0 | 2.7269 | 0 | 0 |
| BBOF1 | 63 kDa | Q3V079 | 2.7269 | 0 | 0 |
| LCA5L | 81 kDa | Q8C0X0 | 2.7269 | 0 | 0 |
| MAPRE3 | 32 kDa | Q6PER3 | 2.7269 | 0 | 0 |
| SMCP | 15 kDa | P15265 | 2.7269 | 0 | 0 |
| C1RA | 80 kDa | Q8CG16 | 1.8179 | 8.8336 | 0 |
| CCDC96 | 67 kDa | Q9CR92 | 1.8179 | 0 | 0 |
| CENPV | 28 kDa | Q9CXS4 | 1.8179 | 2.2084 | 0 |
| LRRC23 | 39 kDa | O35125 | 1.8179 | 0 | 0 |
| KIF3B | 85 kDa | Q61771 | 1.8179 | 0 | 0 |
| RUVBL1 | 50 kDa | P60122 | 1.8179 | 0 | 0 |
| GAS8 | 56 kDa | Q60779 | 1.8179 | 0 | 0 |
| MAPRE1 | 30 kDa | Q61166 | 1.8179 | 0 | 0 |
| LRRC34 | 47 kDa | Q9DAM1 | 1.8179 | 0 | 0 |
| TEPP | 25 kDa | Q6IMH0 | 1.8179 | 0 | 0 |
| CLIP1 | 156 kDa | Q922J3 | 1.8179 | 0 | 0 |
| SPAG8 | 51 kDa | Q3V0Q6 | 1.8179 | 0 | 0 |
| TEX55 | 44 kDa | A6X8Z9 | 1.8179 | 0 | 0 |
| NECTIN3 | 61 kDa | Q9JLB9 | 1.8179 | 0 | 0 |
| TEX47 | 29 kDa | Q9D5W8 | 1.8179 | 0 | 0 |
| HNRNPU | 88 kDa | Q8VEK3 | 0.90896 | 3.3126 | 0 |
| TPP2 | 140 kDa | Q64514 | 0 | 4.4168 | 0 |

**H**

| Protein | MW | Accession Number | Quantitative Value (Normalized Total Spectra) IP: αHA ΔR-Homo | KO |
|---|---|---|---|---|
| GSTM6 | 26 kDa | O35660 | 9.7135 | 0 |
| PGAM1 | 29 kDa | Q9DBJ1 | 8.6342 | 0 |
| MYH11 | 227 kDa | O08638 | 10.793 | 0 |
| VDAC1 | 32 kDa | Q60932 | 7.5549 | 0 |
| ACYP1 | 11 kDa | P56376 | 7.5549 | 0 |
| CANX | 67 kDa | P35564 | 3.2378 | 0 |
| ATP1B1 | 35 kDa | P14094 | 2.1585 | 0 |
| PDCD5 | 14 kDa | P56812 | 7.5549 | 0 |
| TPM2 | 33 kDa | P58774 | 3.2378 | 0 |
| MSTO1 | 61 kDa | Q2YDW2 | 2.1585 | 0 |
| DDX46 | 117 kDa | Q569Z5 | 2.1585 | 0 |
| COPB2 | 102 kDa | O55029 | 2.1585 | 0 |
| PGM3 | 59 kDa | Q9CYR6 | 2.1585 | 0 |
| NDUFV2 | 27 kDa | Q9D6J6 | 2.1585 | 0 |
| JUP | 82 kDa | Q02257 | 4.3171 | 0 |
| COPG2 | 98 kDa | Q9QXK3 | 2.1585 | 0 |
| DPP3 | 83 kDa | Q99KK7 | 3.2378 | 0 |
| PREP | 81 kDa | Q9QUR6 | 3.2378 | 0 |
| TRIM41 | 72 kDa | Q5NCC3 | 3.2378 | 0 |
| SRSF3 | 19 kDa | P84104 | 2.1585 | 0 |
| MPI | 47 kDa | Q924M7 | 3.2378 | 0 |
| ATP5PB | 29 kDa | Q9CQQ7 | 2.1585 | 0 |
| SOD1 | 16 kDa | P08228 | 2.1585 | 0 |
| DHX15 | 91 kDa | O35286 | 2.1585 | 0 |
| CKS2 | 10 kDa | P56390 | 2.1585 | 0 |

**I**

| Protein | MW | Accession Number | Quantitative Value (Normalized Total Spectra) IP: αTRIM41 WT | ΔR-Homo | KO | IP: αHA ΔR-Homo | KO |
|---|---|---|---|---|---|---|---|
| CENPV | 28 kDa | Q9CXS4 | 2.2084 | 0 | 0 | 1.8179 | 0 |
| TRIM41 | 72 kDa | Q5NCC3 | 1.1042 | 3.2378 | 0 | 22.724 | 0 |
| C1RA | 80 kDa | Q8CG16 | 8.8336 | 0 | 0 | 1.8179 | 0 |
| HSP90AB1 | 83 kDa | P11499 | 1.1042 | 0 | 0 | 2.7269 | 0 |
| HNRNPU | 88 kDa | Q8VEK3 | 3.3126 | 0 | 0 | 0.90896 | 0 |
| CCDC40 | 137 kDa | Q8BI79 | 1.1042 | 0 | 0 | 6.3627 | 0 |
| TPP2 | 140 kDa | Q64514 | 4.4168 | 0 | 0 | 0 | 0 |
